# Conditional Guide RNAs: Programmable Conditional Regulation of CRISPR/Cas Function in Bacteria via Dynamic RNA Nanotechnology

**DOI:** 10.1101/525857

**Authors:** Mikhail H. Hanewich-Hollatz, Zhewei Chen, Jining Huang, Lisa M. Hochrein, Niles A. Pierce

## Abstract

A guide RNA (gRNA) directs the function of a CRISPR protein effector to a target gene of choice, providing a versatile programmable platform for engineering diverse modes of synthetic regulation (edit, silence, induce, bind). However, the fact that gRNAs are constitutively active places limitations on the ability to confine gRNA activity to a desired location and time. To achieve programmable control over the scope of gRNA activity, here we apply principles from dynamic RNA nanotechnology to engineer conditional guide RNAs (cgRNAs) whose activity is dependent on the presence or absence of an RNA trigger. These cgRNAs are programmable at two levels, with the trigger-binding sequence controlling the scope of the effector activity and the target-binding sequence determining the subject of the effector activity. We demonstrate molecular mechanisms for both constitutively active cgRNAs that are conditionally inactivated by an RNA trigger (ON→OFF logic) and constitutively inactive cgRNAs that are conditionally activated by an RNA trigger (OFF→ON logic). For each mechanism, automated sequence design is performed using the reaction pathway designer within NUPACK to design an orthogonal library of three cgRNAs that respond to different RNA triggers. In *E. coli* expressing cgRNAs, triggers, and silencing dCas9 as the protein effector, we observe programmable conditional gene silencing with a median dynamic range of ≈6-fold for an ON→OFF “terminator switch” mechanism, ≈15-fold for an ON→OFF “splinted switch” mechanism, and ≈3.6-fold for an OFF→ON “toehold switch” mechanism; the median crosstalk within each cgRNA library is <2%, <2%, and ≈20% for the three mechanisms. By providing programmable control over both the scope and target of protein effector function, cgRNA regulators offer a promising platform for synthetic biology.

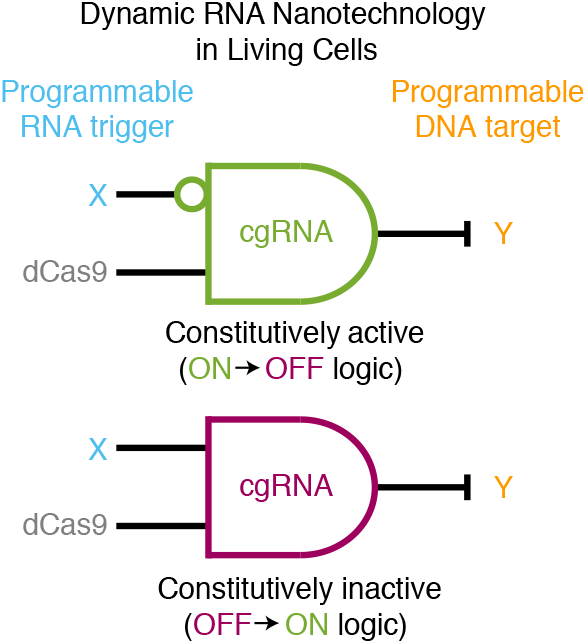

## INTRODUCTION

Dynamic RNA nanotechnology holds great promise as a paradigm for introducing synthetic regulatory links into living cells and organisms. We envision small conditional RNAs (scRNAs) that, upon detection of a programmable nucleic acid input, change conformation to produce a programmable output that up-regulates or down-regulates the activity of a biological pathway. In this scenario, the input controls the scope of regulation and the output controls the target of regulation, with the scRNA performing signal transduction to create a logical link between the two.^1,2^ Any pathway that recognizes RNA is a potential candidate for conditional regulation by scRNAs (e.g., RNA interference, RNase H, PKR, RIG-1); the CRISPR/Cas pathway is a particularly attractive candidate because of its functional versatility, high regulatory dynamic range, and portability between species.^3–5^

The repurposing of RNA-guided CRISPR effectors through development of modified guide RNAs (gRNAs) and CRISPR-associated (Cas) proteins has yielded a suite of powerful tools for biological research and synthetic biology. Precision genome editing has been achieved in a variety of organisms using gRNAs to direct the nuclease activity of Cas9 and Cas12a (Cpf1) to a target gene of choice.^3,6–8^ Mutation of the nuclease domains to produce a catalytically dead Cas9 (dCas9) has enabled silencing of genetic expression via inhibition of transcriptional elongation,^4,9^ or induction of genetic expression using dCas9 fusions that incorporate transcriptional regulatory domains.^5^ Other dCas9 fusions have mediated target-binding to enable visualization of genomic loci,^10,11^ epigenetic modification,^12^ and single-base editing at a specific genomic locus.^3, 13^ Hence, gRNA:effector complexes combine the benefits of the rich functional vocabulary of the protein effector (edit, silence, induce, bind) and the programmability of the gRNA in targeting effector activity to a gene of choice.

Because gRNAs are constitutively active, additional measures are needed to restrict effector activity to a desired location and time. Temporal control can be achieved by small-molecule induction of gRNAs^14,15^ or Cas9,^16^ but this comes with limitations in terms of multiplexing and spatial control. Spatiotemporal control has been achieved by regulation of Cas9 via photoactivation^17^ or via tissue-specific promoters^18, 19^ or microRNAs,^20^ which comes with the unwelcome restriction that all gRNAs are subject to the same regulatory scope. Systematic mapping of the structure and sequence properties of functional gRNAs has revealed that Cas9 activity is tolerant to significant modifications the standard gRNA structure,^21,22^ facilitiating introduction of auxiliary domains that enable conditional control of gRNA activity via structural changes induced by small-molecules,^23,24^ protein-bound RNAs,^25^ nucleases,^26^ or nuclease-recruiting DNAs.^26^ Alternatively, the activity of standard gRNAs has been modulated by antisense RNAs^27^ or by photolysis of antisense DNAs incorporating photocleavable groups.^28^

Here, pursuing the scRNA paradigm of programmable conditional regulation based on dynamic RNA nanotechnology, and leveraging information on tolerated modifications of standard gRNA structure, we set out to engineer conditional guide RNAs (cgRNAs) that change conformation in response to an RNA trigger X to conditionally direct the function of dCas9 to a target gene Y. Unlike a standard gRNA, a cgRNA is programmable at two levels, with the trigger-binding sequence controlling the scope of cgRNA activity and the target-binding sequence determining the subject of effector activity. Functionally, the cgRNA must perform sequence transduction between X and Y as well as shape transduction between active/inactive conformations. cgRNA activity can be engineered to toggle either OFF→ON (as was recently demonstrated by Siu and Chen^29^) or ON→OFF in response to a cognate RNA trigger X; this conditional control can be exerted over dCas9 variants that either edit, silence, induce, or bind the target Y, emphasizing the broad functional potential available via interplay between cgRNA logic and protein effector function (Figure 1a).

**Figure 1.**
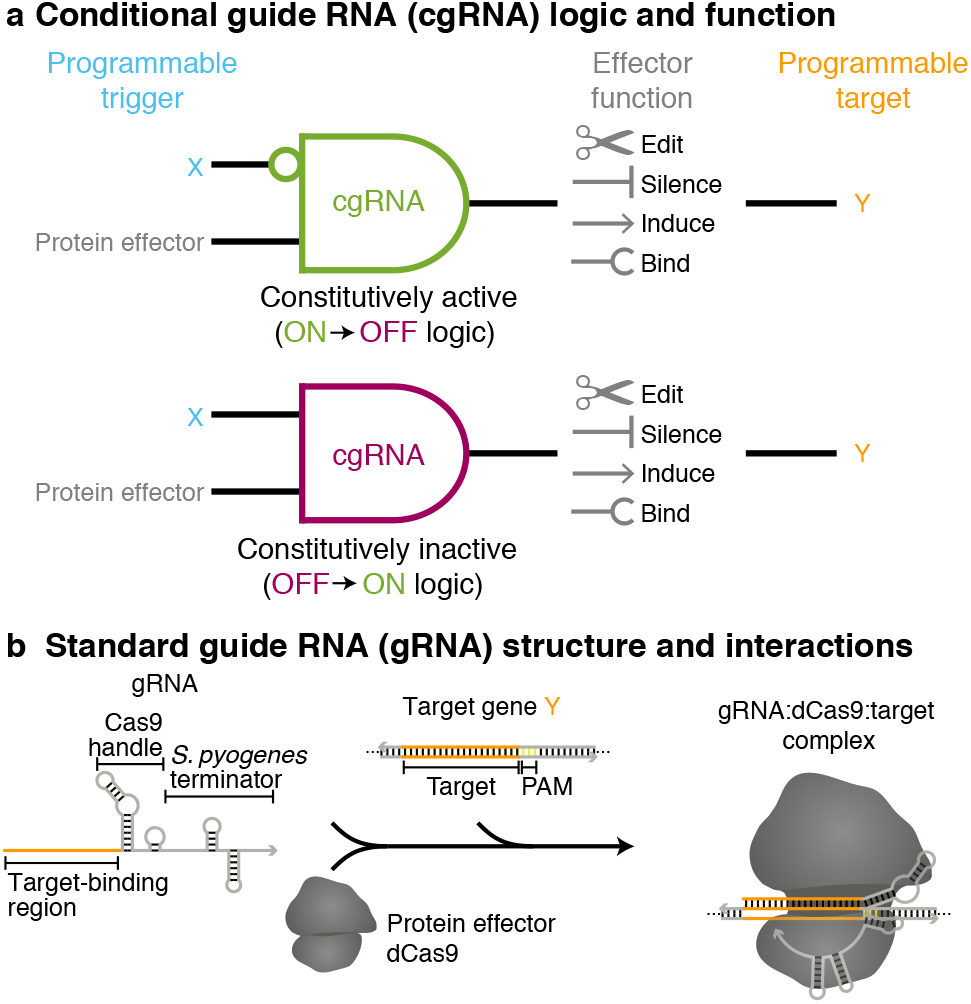
Programmable regulators. (a) A conditional guide RNA (cgRNA) changes conformation in response to a programmable trigger X to conditionally direct the activity of a protein effector to a programmable target Y. Top: a constitutively active cgRNA is conditionally inactivated by X (ON→OFF logic). Bottom: a constitutively inactive cgRNA is conditionally activated by X (OFF→ON logic). (b) A standard guide RNA (gRNA) is constitutively active, directing the function of protein effector dCas9 to a target gene Y; different dCas9 variants implement different functions (edit, silence, induce, bind).

## RESULTS AND DISCUSSION

### Constitutively Active Terminator Switch cgRNA Mechanism (ON→OFF logic)

As a starting point, consider the constitutively active “terminator switch” cgRNA mechanism of Figure 2a that is conditionally inactivated by RNA trigger X (ON→OFF logic). Compared to a standard gRNA (Figure 1b), the cgRNA has a modified terminator region with an extended loop and rationally designed sequence domains “d-e-f”. Hybridization of the RNA trigger X to these modified domains is intended to form a structure incompatible with cgRNA mediation of dCas9 function. We validated the cgRNA mechanism in *vivo* in *E. coli* expressing silencing dCas9^4^ as the protein effector and a fluorescent protein reporter (mRFP) as the target gene Y. A cell line expressing the cgRNA exhibits low fluorescence (ON state) while a cell line expressing both the cgRNA and the cognate RNA trigger exhibit high fluorescence (OFF state), achieving a conditional response of an order of magnitude (Figure 2b). With the terminator switch mechanism, the sequences of the RNA trigger X and the silencing target Y are fully independent, with the cgRNA mediating allosteric regulation – the trigger down-regulates cgRNA:dCas9 function not by sequestering the target-binding region (orange in Figure 2a), but by hybridizing to the distal trigger-binding region (blue). Ideally, a cgRNA would have a strong ON state with activity equivalent to a standard gRNA and a clean OFF state with no activity. Testing control lines expressing a standard gRNA targeting Y (ideal ON state) or a no-target gRNA that lacks the target-binding region (ideal OFF state), we observe a dynamic range of two orders of magnitude, demonstrating room for future improvement of both the ON state and the OFF state for this cgRNA mechanism. To test programmability, we used NUPACK^30,31^ to design a library of three orthogonal cgRNA/trigger pairs (Figure 2c), achieving a median ≈6-fold conditional ON→OFF response to expression of the cognate trigger (left) and median crosstalk below 2% between noncognate cgRNA/trigger combinations (right).

**Figure 2.**
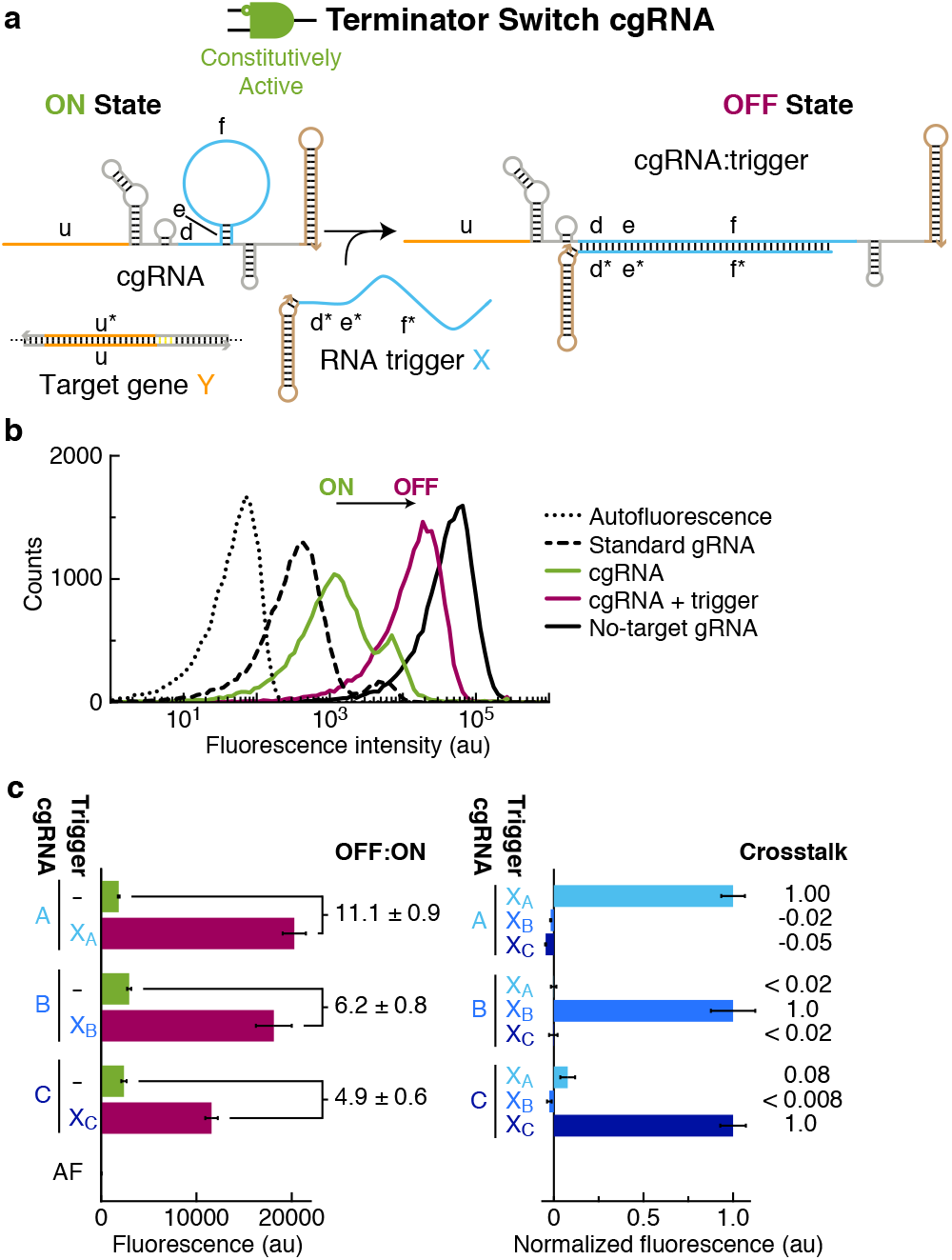
Conditional gene silencing using constitutively active terminator switch cgRNA in bacteria (ON→OFF logic). (a) Mechanism schematic. The constitutively active cgRNA is inactivated by hybridization of RNA trigger X. Rational sequence design of cgRNA terminator region (domains “d-e-f’ comprising 6 nt linker, 4 nt stem, 30 nt loop) and complementary trigger region (domains “f*-e*-d*”). (b) Expression of RNA trigger X (40 nt + synthetic terminator) toggles the cgRNA from ON→OFF, leading to an increase in fluorescence. Single-cell fluorescence intensities via flow cytometry. Induced expression (aTc) of silencing dCas9 and constitutive expression of mRFP target gene Y and either: standard gRNA (ideal ON state), cgRNA(ON state), cgRNA + RNA trigger X (OFF state; trigger expression is IPTG-induced), no-target gRNA that lacks target-binding region (ideal OFF state). Autofluorescence (AF): cells with no mRFP. (c) Programmable conditional regulation using 3 orthogonal cgRNAs (A, B, C). Left: ON→OFF conditional response to cognate trigger (OFF:ON ratio = [OFF–AF]/[ON–AF]). Right: crosstalk between non-cognate cgRNA/trigger pairs ([non-cognate trigger – no trigger]/[cognate trigger – no trigger]). Fluorescence via flow cytometry: mean ± estimated standard error of the mean (with uncertainty propagation) of median single-cell fluorescence over 20,000 cells, *N* = 3 replicate wells.

### Constitutively Active Splinted Switch cgRNA Mechanism (ON→OFF logic)

To test an alternative approach to implementing this ON→OFF conditional logic, we next tested a constitutively active “splinted switch” cgRNA mechanism (Figure 3a) that has extended loops in both the Cas9 handle (domain “d”) and terminator (domain “e”). Hybridization of RNA trigger X to both loops is intended to form a splint that is structurally incompatible with cgRNA mediation of dCas9 function. In *E. coli* expressing silencing dCas9 and a fluorescent protein reporter (sfGFP) as the target gene Y, the splinted switch achieves an order of magnitude ON→OFF conditional response to expression of RNA trigger X (Figure 3b). Notably, the ON state is similar to that of a standard gRNA, implying that dCas9 is more tolerant of the splinted switch modifications to the handle and terminator loops than it was to the terminator switch modifications to the terminator linker/stem/loop. However, these gains in ON state are partially offset by losses in the cleanliness of the OFF state, which falls short of the no-target gRNA control by an order of magnitude. Examining a library of three orthogonal splinted switch cgRNA/trigger pairs designed using NUPACK (Figure 3c), we observe a median ≈15-fold ON→OFF conditional response to expression of the cognate trigger and median crosstalk below 2% between non-cognate cgRNA/trigger combinations. As with the terminator switch mechanism, splinted switch cgRNAs are allosteric regulators – the trigger down-regulates cgRNA:dCas9 function by hybridizing to extended loops (blue in Figure 3a) distal to the target-binding region (orange). The resulting full sequence independence between RNA trigger X and target gene Y provides the flexibility for X to control regulatory scope independent of the choice of Y.

**Figure 3.**
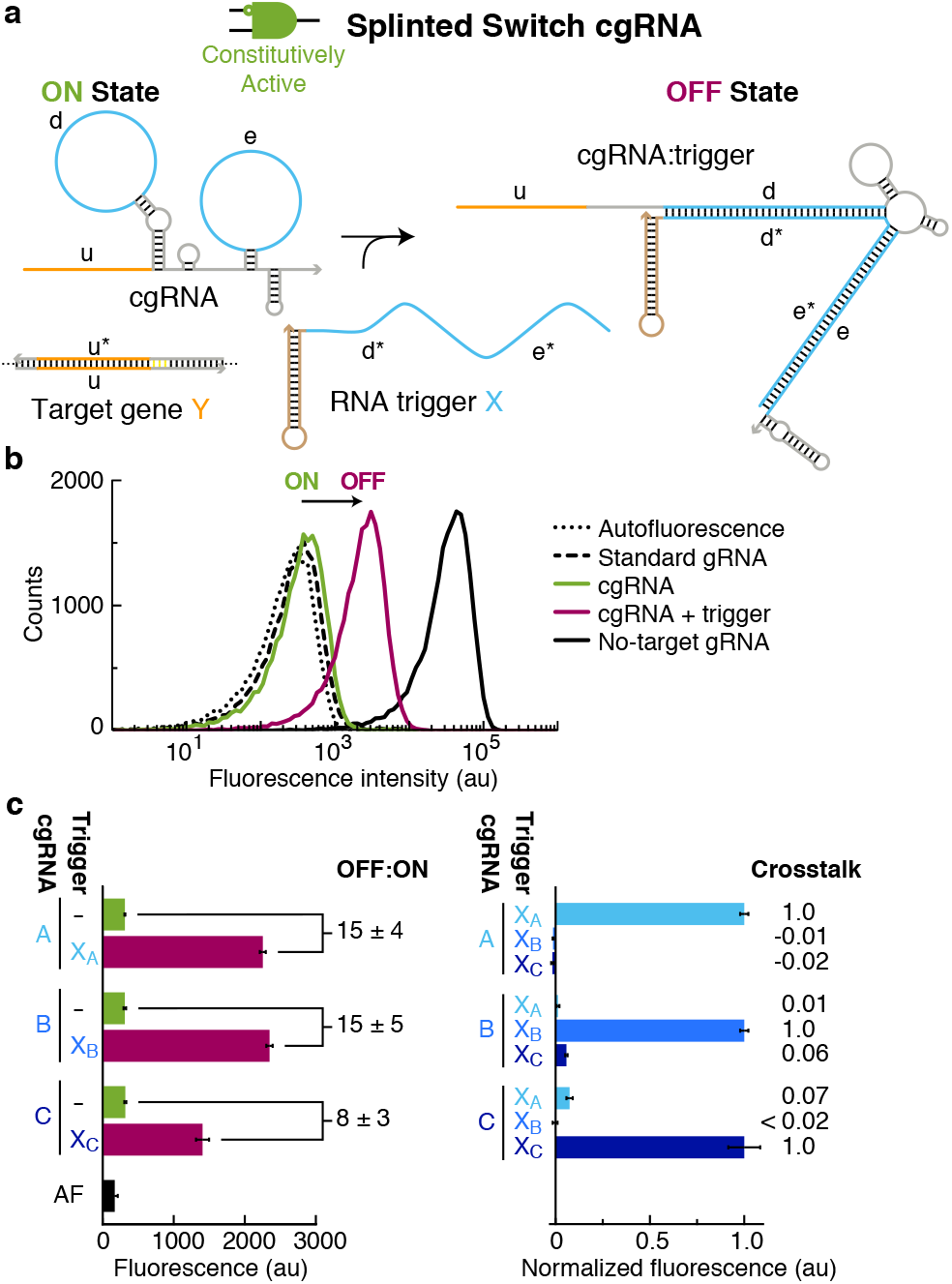
Conditional gene silencing using constitutively active splinted switch cgRNAs in bacteria (ON→OFF logic). (a) Mechanism schematic. The constitutively active cgRNA is inactivated by hybridization of RNA trigger X. Rational sequence design of the 35 nt Cas9 handle loop (domain “d”) and an extended 35 nt terminator hairpin loop (domain “e”). (b) Expression of RNA trigger X (70 nt + synthetic terminator) toggles the cgRNA from ON→OFF, leading to an increase in fluorescence. Single-cell fluorescence intensities via flow cytometry. Induced expression (aTc) of silencing dCas9 and constitutive expression of sfGFP target gene Y and either: standard gRNA (ideal ON state), cgRNA(ON state), cgRNA + RNA trigger X (OFF state), notarget gRNA that lacks target-binding region (ideal OFF state). Autofluorescence (AF): cells with no sfGFP. (c) Programmable conditional regulation using 3 orthogonal cgRNAs (A, B, C). Left: ON→OFF conditional response to cognate trigger (OFF:ON ratio = [OFF–AF]/[ON–AF]). Right: crosstalk between non-cognate cgRNA/trigger pairs ([non-cognate trigger – no trigger]/[cognate trigger – no trigger]). Fluorescence via flow cytometry: mean ± estimated standard error of the mean (with uncertainty propagation) of median single-cell fluorescence over 20,000 cells, *N* = 3 replicate wells.

### Constitutively Inactive Toehold Switch cgRNA Mechanism (OFF→ON logic)

To reverse the conditional logic, we then tested a constitutively inactive “toehold switch” cgRNA mechanism (Figure 4a) that is conditionally activated by RNA trigger X (OFF:ON logic). The target-binding region of the cgRNA (domain “u”) is initially sequestered by a 5’ extension to inhibit recognition of target gene Y; hybridization of trigger X to this extension is intended to desequester the target-binding region and enable cgRNA direction of dCas9 function to target gene Y. In *E. coli* expressing silencing dCas9 and a fluorescent protein reporter (mRFP) as the target gene Y, the toehold switch cgRNA achieves less than an order of magnitude OFF:ON conditional response to expression of RNA trigger X (Figure 4b). In this case, the OFF state is degraded by an order of magnitude relative to the ideal OFF state (no-target gRNA control) and the ON state is degraded by an order of magnitude relative to the ideal ON state (standard gRNA control). For a library of three orthogonal toehold switch cgRNA/trigger pairs designed using NUPACK (Figure 4c), we observe a median ≈3.6-fold OFF:ON conditional response to expression of the cognate trigger and median crosstalk of ≈20% between non-cognate cgRNA/trigger combinations. Recently, Siu and Chen demonstrated a median ≈6.6-fold OFF:ON conditional response using toehold switch cgRNAs with subtly different structural details in the sequestration of the target-binding region.^29^ Unlike the terminator switch and splinted switch mechanisms for ON→OFF logic, toehold switch cgRNAs for OFF:ON logic are not allosteric as the cgRNA initially down-regulates cgRNA:dCas9 function by sequestering the target-binding region (orange domain “u” in Figure 4a) with a portion of the trigger-binding region (orange domain “u*”). As a result, the toehold switch cgRNAs offer only partial sequence independence between the trigger X and the target gene Y (“u” is a subsequence of both X and Y). This partial sequence dependence is not necessarily limiting for synthetic biology applications where the trigger can be rationally designed and expressed exogenously, but does pose a limitation in situations where X and Y are both endogenous sequences.

**Figure 4.**
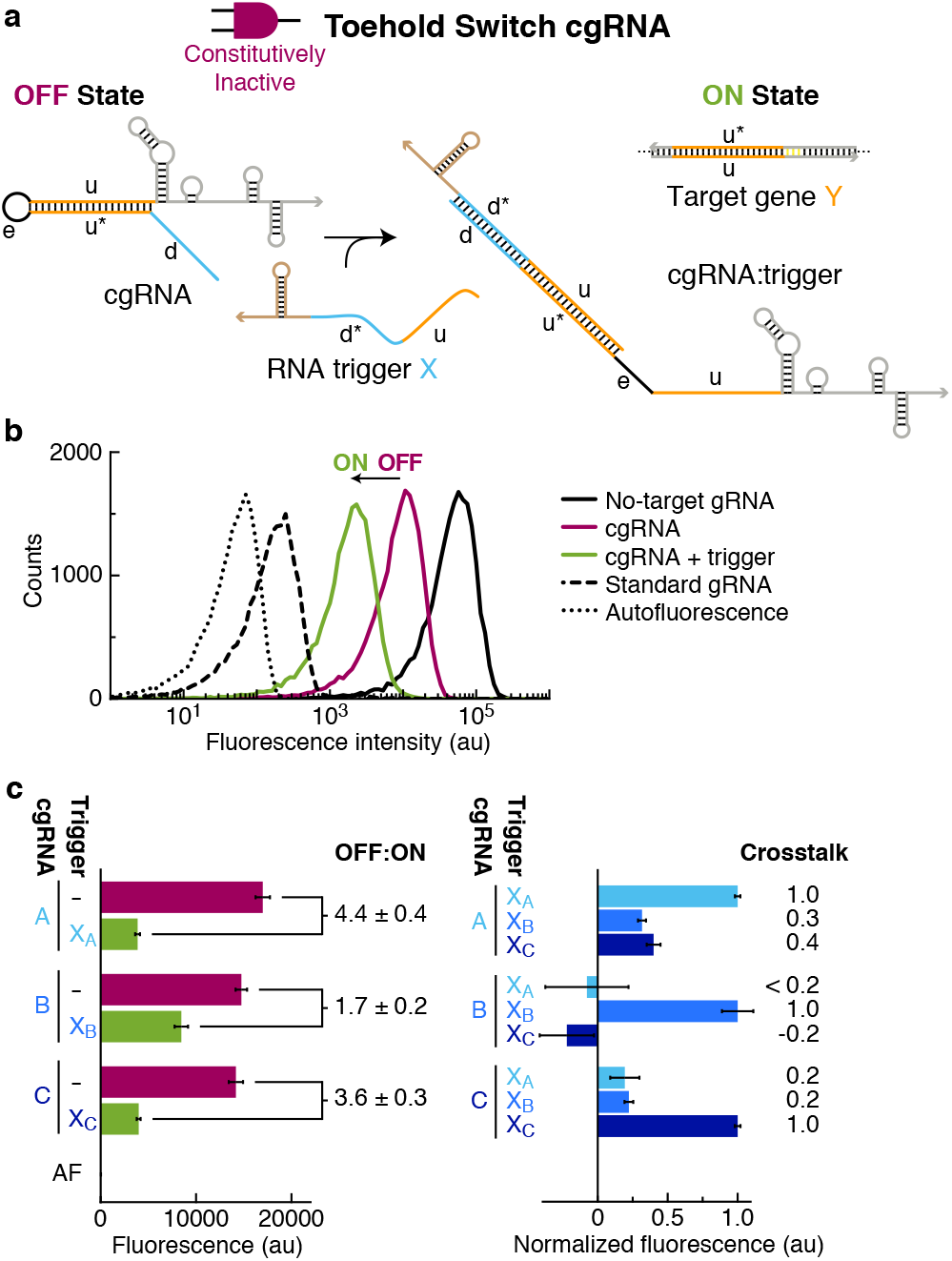
Conditional gene silencing using constitutively inactive toehold switch cgRNAs in bacteria (OFF→ON logic). (a) Mechanism schematic. The constitutively inactive cgRNA is activated by hybridization of RNA trigger X. Rational sequence design of the toehold (domain “d”; 15 nt) and loop (domain “e”; 8 nt) flanking the sequestration domain “u*” (20 nt). (b) Expression of RNA trigger X (35 nt + synthetic terminator) toggles the cgRNA from OFF→ON, leading to a decrease in fluorescence. Single-cell fluorescence intensities via flow cytometry. Induced expression (aTc) of silencing dCas9 and constitutive expression of mRFP target gene Y and either: no-target gRNA that lacks target-binding region (ideal OFF state), cgRNA (OFF state), cgRNA + RNA trigger X (ON state), standard gRNA (ideal ON state). Autofluorescence (AF): cells with no mRFP. (c) Programmable conditional regulation using 3 orthogonal cgRNAs (A, B, C). Left: OFF→ON conditional response to cognate trigger (OFF:ON ratio = [OFF–AF]/[ON–AF]). Right: crosstalk between non-cognate cgRNA/trigger pairs ([non-cognate trigger – no trigger]/[cognate trigger – no trigger]). Fluorescence via flow cytometry: mean ± estimated standard error of the mean (with uncertainty propagation) of median single-cell fluorescence over 20,000 cells, *N* = 3 replicate wells.

### Computational Sequence Design of Libraries of Orthogonal cgRNAs using NUPACK

For each cgRNA mechanism (Figures 2–4), sequence design was performed using the reaction pathway designer within NUPACK.^30,31^ Following Wolfe et al.,^31^ sequence design was formulated as a multistate optimization problem using target test tubes to represent reactant and product states of cgRNA/trigger hybridization, as well as to model crosstalk between orthogonal cgRNAs (Figure 5a). Each reactants tube (Step 0) and products tube (Step 1) contains a set of desired “on-target” complexes (each with a target secondary structure and target concentration) corresponding to the on-pathway hybridization products for a given step, and a set of undesired “off-target” complexes (each with a target concentration of 0 nM) corresponding to on-pathway reactants and off-pathway hybridization crosstalk for a given step. Hence, these elementary step tubes are designed for full conversion of cognate reactants into cognate products and against local hybridization crosstalk between these same reactants. To simultaneously design *N* orthogonal systems, elementary step tubes are specified for each system (Figure 5a; left). Furthermore, to design against off-pathway interactions between systems, a single global crosstalk tube is specified (Figure 5a; right). In the global crosstalk tube, the on-target complexes correspond to all reactive species generated during all elementary steps (*m* = 0,1) for all systems (*n* = 1,…,*N*); the off-target complexes correspond to non-cognate interactions between these reactive species. Crucially, the global crosstalk tube ensemble omits the cognate products that the reactive species are intended to form (they appear as neither on-targets nor off-targets). Hence, all reactive species in the global crosstalk tube are forced to either perform no reaction (remaining as desired on-targets) or undergo a crosstalk reaction (forming undesired off-targets), providing the basis for minimization of global crosstalk during sequence optimization.

**Figure 5.**
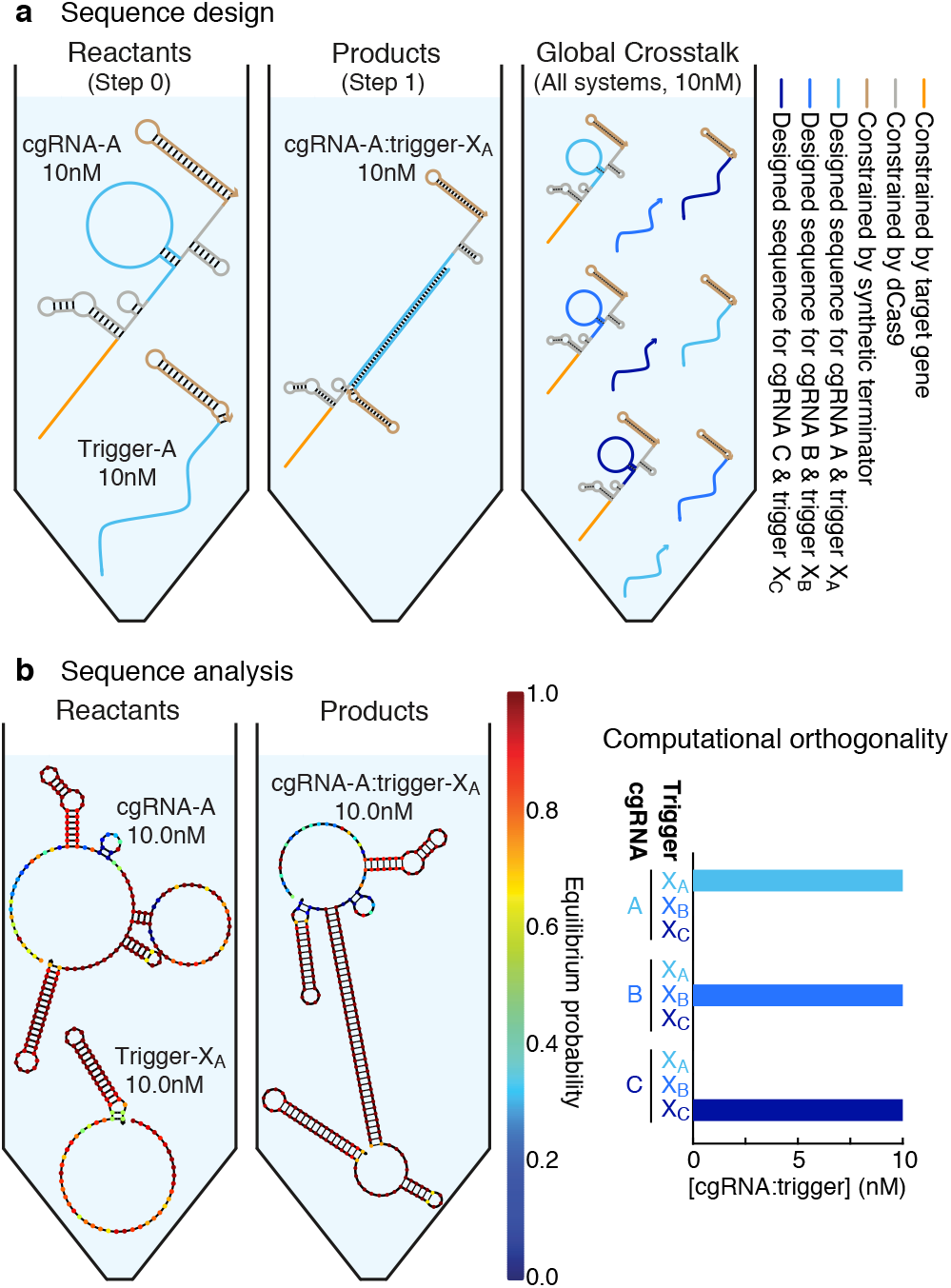
Computational cgRNA sequence design using NUPACK.^30,31^. (a) Target test tubes for design of 3 orthogonal cgR-NAs A, B, C (terminator switch mechanism for Figure 2). Left: Elementary step tubes. Reactants tube (Step 0): cgRNA and trigger. Products tube (Step 1): cgRNA:trigger. Each target test tube contains a set of desired “on-target” complexes (each with the depicted target secondary structure and a target concentration of 10 nM) corresponding to the on-pathway hybridization products for a given step and a set of undesired “off-target” complexes (all complexes of up to 2 strands, each with a target concentration of 0 nM; not depicted) corresponding to on-pathway reactants and off-pathway hybridization crosstalk for a given step. To design 3 orthogonal systems, there are two elementary step tubes for each system A, B, C. Right: Global crosstalk tube. Contains the depicted on-target complexes corresponding to reactive species generated during Steps 0 and 1 (each with the depicted target secondary structure and a target concentration of 10 nM) as well as off-target complexes (all complexes of up to 2 strands, each with a target concentration of 0 nM; not depicted) corresponding to off-pathway interactions between these reactive species. To design 3 orthogonal systems, the global crosstalk tube contains a set of on-targets and off-targets for each system A, B, C. (b) Analysis of design quality.^30, 32^ Left: Tubes depict the target structure and predicted concentration for each on-target complex with nucleotides shaded to indicate the probability of adopting the depicted base-pairing state at equilibrium. For this design, all on-targets are predicted to form with quantitative yield at the 10 nM target concentration but some nucleotides have unwanted base-pairing interactions (nucleotides not shaded dark red). Right: Computational orthogonality study. Predicted equilibrium concentration of each cgRNA:trigger complex for the 3 orthogonal systems of Figure 2 (one cgRNA species and one RNA trigger species per tube). RNA at 37°C in 1M Na^+^.^33^

Sequence design is performed subject to complementarity constraints inherent to the reaction pathway (Figure 2a; domain “d” complementary to “d*”, etc.), as well as to biological sequence constraints imposed by the the silencing target Y (mRFP or sfGFP), the protein effector (dCas9), or the synthetic terminator; see the constraint shading in Figure 5a). The sequence is optimized by reducing the ensemble defect quantifying the average fraction of incorrectly paired nucleotides over the multi-tube ensemble.^31,34,35^ Within the ensemble defect, defect weights were applied to prioritize design effort.^31^ Optimization of the ensemble defect implements both a positive design paradigm, explicitly designing for on-pathway elementary steps, and a negative-design paradigm, explicitly designing against off-pathway crosstalk.^31^

Figure 5b displays the Reactants and Products tubes for a completed sequence design (cgRNAs of Figure 2). For cgRNA A (left panel), on-target complexes are predicted to form with quantitative yield at the target concentrations, but with some unintended base-pairing (nucleotides not shaded dark red). These structural defects within the ensemble of on-target complexes reflect the real-world challenges of designing a cgRNA that satisfies biological sequence constraints, changes conformation in response to a cognate RNA trigger, and operates orthogonally to a library of other cgRNAs. For the corresponding library of orthogonal cgRNAs (A, B, C), each cgRNA is predicted to interact appreciably only with its cognate RNA trigger (right panel).

### Conceptual Opportunities for cgRNA-Enabled Biological Research Tools, Therapeutics, and Synthetic Biology

The ability to rationally design cgRNAs suggests a conceptual framework for enabling biologists to exert spatiotemporal control over regulatory perturbations in living organisms using CRISPR/Cas technology. In principle, Cas activity could be restricted to a desired cell type, tissue, or organ by selecting an endogenous RNA trigger X with the desired spatial and temporal expression profile. To shift conditional regulation to a different tissue or developmental stage, the cgRNA would be reprogrammed to recognize a different trigger sequence. Signal transduction with cgRNAs would also have attractive therapeutic potential, with trigger X as a programmable disease marker and target Y as an independent programmable therapeutic target, enabling selective treatment of diseased cells. Synthetic biology provides another attractive arena for use of cgRNAs. Traditional synthetic biology regulators have relied on protein:protein and protein:DNA interactions mined from existing genomes, placing limits on scalability due to crosstalk and the limited number of available regulators. cgRNA regulators offer a promising platform for scalable synthetic biology.

### Comparison of cgRNAs to other scRNAs

It is interesting to compare the present work engineering cgRNAs (a particular class of scRNAs with notable properties) to the scRNAs we previously engineered toward the goal of conditional RNA interference (RNAi).^1,2^ In both cases, the scRNAs are intended to perform signal transduction between detection of a programmable RNA input and production of a biologically active programmable output. In the case of conditional RNAi, the scRNAs detect an mRNA input X and interact to produce a substrate that is processed by Dicer to produce an siRNA output targeting independent silencing target mRNA Y for destruction. Because Dicer substrates are structurally simple, comprising predominantly a duplex containing the target-binding sequence,^36^ signal transduction between X and Y and inactive/active states is performed by scRNAs upstream of formation of the biologically active Dicer substrate. For example, the simplest mechanism we have devised to date involves a dimer scRNA that conditionally generates a monomer Dicer substrate anti-Y upon detection of mRNA X.^1,2^ By contrast, the standard gRNAs that serve as substrates for Cas9 protein effectors are not only structurally more complex than Dicer substrates (involving multiple duplexes, loops, and tails), but Cas9 also appears to be more permissive of modifications to the standard structure, providing hooks for engineering programmable conditional regulation. As a result, it is possible to do signal transduction between X and Y and inactive/active states (for either ON→OFF or OFF→ON logic) all within a single cgRNA (i.e., a single monomer scRNA). A benefit of this mechanistic simplicity is that monomer cgRNAs can be readily expressed, while expression of well-formed multimer scRNAs such as those developed for conditional Dicer substrate formation appears more challenging, possibly necessitating delivery with chemical reagents.

## CONCLUSIONS

The present work represents only a first step toward our longterm goal of engineering programmable conditional regulators that function robustly in living organisms. Here, we made progress on several fronts: 1) demonstrating cgRNA mechanisms *in vivo* in bacteria for both constitutively active cgRNAs that are inactivated by a cognate RNA trigger (ON→OFF logic) and constitutively inactive cgRNAs that are conditionally activated by a cognate RNA trigger (OFF→ON logic), 2) demonstrating the applicability and versatility of dynamic RNA nanotechnology in living cells, and 3) demonstrating automated sequence design using NUPACK to engineer libraries of orthogonal cgRNA/trigger pairs.

In order to develop cgRNAs into a versatile platform for biological research, a number of major improvements are needed. First, standard gRNAs routinely achieve two orders of magnitude in regulatory dynamic range,^4^ and it is desirable to engineer improved cgRNA mechanisms that exploit this full dynamic range. Toward this end, further understanding of the structure/function relationships between cgRNAs, triggers, and Cas effectors are needed to ascertain how to robustly achieve both a strong ON state and a clean OFF state depending on the presence/absence of the cognate trigger. Second, to enable tissue-selective regulation in living organisms, it is critical that cgRNAs are able to efficiently detect a trigger that is a subsequence of a longer endogenous RNA (e.g., a subsequence of an mRNA). Detection of a subsequence of a full-length mRNA poses significant additional challenges relative to detection of a short RNA trigger,^2,29^ increasing the degree of difficulty in achieving a conditional response that exploits the full dynamic range. Third, in common with the terminator switch and splinted switch mechanisms studied here (but unlike the toehold switch mechanisms studied here and elsewhere^29^), it is important that cgRNA regulators be allosteric, so that the sequence of the target gene Y places no restriction on the sequence of the RNA trigger X, enabling independent control over the regulatory scope (using X) and the regulatory target (using Y). Significant effort and innovation are needed to achieve these goals and develop cgRNAs that operate as plug-and-play programmable conditional regulators within endogenous biological circuits in living organisms.

## METHODS SUMMARY

For each mechanism, orthogonal cgRNA/trigger pairs were designed using the reaction pathway engineering tools within NUPACK (nupack.org).^30,31^ Control gRNA and cgRNA/trigger constructs were generated by inserting sequence modifications into a previously described pgRNA-bacteria vector.^4^ A modified *E. coli* MG1655 strain containing genomically incorporated mRFP and sfGFP^4^ was used for all fluorescence assays. Sequence verified strains were grown overnight in EZ-RDM (Teknova) containing carbenicillin and chloramphenicol and seeded at 100 × dilution in fresh medium and grown to midlog phase (≈4 h), then further diluted ≈100× to normalize cell density with fresh medium containing aTc for induction of Cas9 expression (and 5mM IPTG for terminator switch experiments only). Induced cells were grown at 37 °C with continuous shaking for 12 h, with end-point fluorescence measured via flow cytometry (20,000 live cell counts per well).

## ASSOCIATED CONTENT

### Supporting Information

Methods, sequences, plasmids, schematics, flow cytometry replicates. This material is available free of charge via the Internet at http://pubs.acs.org.

### Notes

The authors declare competing financial interests in the form of pending patents.

## ACKNOWLEDGEMENTS

We thank S. Qi for the gift of plasmids and the gift of *E. coli* expressing mRFP and sfGFP, N.J. Porubsky for assistance with reaction pathway engineering using NUPACK, A. Hou and J. Kishi for performing preliminary studies, and R. Phillips for discussions on allosteric regulation. This work was funded by the Defense Advanced Research Projects Agency (HR0011-17-2-0008; the findings are those of the authors and should not be interpreted as representing the official views or policies of the US Government), by the Caltech Center for Environmental Microbial Interactions (CEMI), by the National Institutes of Health (5T32GM112592), by the Rosen Bioengineering Center at Caltech, by the National Science Foundation Molecular Programming Project (NSF-CCF-1317694), by a Professorial Fellowship at Balliol College (University of Oxford), and by the Eastman Visiting Professorship at the University of Oxford.

