## Supporting Information for "Conditional Guide RNAs: Programmable Conditional Regulation of CRISPR/Cas Function in Bacteria via Dynamic RNA Nanotechnology"

#### Contents

**S1 Methods** **S3**
S1.1 Rational design of libraries of orthogonal cgRNAs using NUPACK . . . S3
S1.1.1 Target test tube specification for terminator switch mechanism . . . S4
S1.1.2 Target test tube specification for splinted switch mechanism . . . S6
S1.1.3 Target test tube specification for toehold switch mechanism . . . S8
S1.2 Plasmid construction and molecular cloning . . . S10
S1.3 Bacterial culture for cgRNA studies . . . S10
S1.4 Flow cytometry . . . S10
S1.5 Cell fluorescence analysis . . . S11
S1.5.1 OFF:ON ratio for constitutively active cgRNAs (ON→OFF logic) . . . S12
S1.5.2 OFF:ON ratio for constitutively inactive cgRNAs (OFF→ON logic) . . . S12
S1.5.3 Crosstalk for orthogonal cgRNAs . . . S12

**S2 Sequences** **S13**
S2.1 Sequences for cgRNAs, triggers, and control gRNAs . . . S13
S2.2 Transcriptional promoter and terminator sequences . . . S14
S2.3 Gene sequences . . . S14

**S3 Plasmids** **S16**
S3.1 Constitutively active terminator switch in *E. coli* . . . S16
S3.2 Constitutively active splinted switch in *E. coli* . . . S19
S3.3 Constitutively inactive toehold switch in *E. coli* . . . S22
S3.4 Plasmid for expression of lacI + dCas9 . . . S25

**S4 Schematics of putative ON and OFF states** **S27**
S4.1 Constitutively active terminator switch cgRNA . . . S27
S4.2 Constitutively active splinted switch cgRNA . . . S27
S4.3 Constitutively inactive toehold switch cgRNA . . . S28

**S5 Flow cytometry replicates** **S29**
S5.1 Constitutively active terminator switch in *E. coli* . . . S29
S5.1.1 ON state, OFF state, and conditional response (cf. Figure 2b) . . . S29
S5.1.2 Orthogonal library studies (cf. Figure 2c) . . . S29
S5.2 Constitutively active splinted switch in *E. coli* . . . S30
S5.2.1 ON state, OFF state, and conditional response (cf. Figure 3b) . . . S30
S5.2.2 Orthogonal library studies (cf. Figure 3c) . . . S30
S5.3 Constitutively inactive toehold switch in *E. coli* . . . S31
S5.3.1 ON state, OFF state, and conditional response (cf. Figure 4b) . . . S31
S5.3.2 Orthogonal library studies (cf. Figure 4c) . . . S31

[†]Division of Biology & Biological Engineering, California Institute of Technology, Pasadena, CA 91125, USA. [‡]Division of Engineering & Applied Science, California Institute of Technology, Pasadena, CA 91125, USA. [¶]Weatherall Institute of Molecular Medicine, University of Oxford, Oxford OX3 9DS, UK. [*]

### List of Figures


S1 Target test tubes for sequence design of orthogonal terminator switch cgRNAs . . . S4
S2 Nucleotide defect weights for sequence design of terminator switch cgRNAs . . . S5
S3 Target test tubes for sequence design of orthogonal splinted switch cgRNAs . . . S6
S4 Nucleotide defect weights for sequence design of splinted switch cgRNAs . . . S7
S5 Target test tubes for sequence design of orthogonal toehold switch cgRNAs . . . S9
S6 Nucleotide defect weights for sequence design of toehold switch cgRNAs . . . S9
S7 Illustration of gates used for flow cytometry analysis of *E. coli*. . . . S11
S8 Example plasmid map for terminator switch . . . S17
S9 Example annotated plasmid sequence for terminator switch . . . S18
S10 Example plasmid map for splinted switch . . . S20
S11 Example annotated plasmid sequence for splinted switch . . . S21
S12 Example plasmid map for toehold switch . . . S23
S13 Example annotated plasmid sequence for toehold switch . . . S24
S14 Plasmid map for pdCas9+lacI . . . S25
S15 Annotated plasmid sequence for expression of lacI + dCas9 . . . S26
S16 Schematics of putative ON and OFF states for terminator switch mechanism . . . S27
S17 Schematics of putative ON and OFF states for splinted switch mechanism . . . S27
S18 Schematics of putative OFF and ON states for toehold switch mechanism . . . S28
S19 Flow cytometry replicates for terminator switch ON state, OFF state, and conditional response in *E. coli* (cf. Figure 2b) . . . S29
S20 Flow cytometry replicates for terminator switch orthogonal response in *E. coli* (cf. Figure 2c) . . . S29
S21 Flow cytometry replicates for splinted switch ON state, OFF state, and conditional response in *E. coli* (cf. Figure 3b) . . . S30
S22 Flow cytometry replicates for splinted switch orthogonal response in *E. coli* (cf. Figure 3c) . . . S30
S23 Flow cytometry replicates for toehold switch ON state, OFF state, and conditional response in *E. coli* (cf. Figure 4b) . . . S31
S24 Flow cytometry replicates for toehold switch orthogonal response in *E. coli* (cf. Figure 4c) . . . S31


### List of Tables


S1 Terminator switch sequences . . . S13
S2 Splinted switch sequences . . . S13
S3 Toehold switch sequences . . . S14
S4 Control gRNA sequences . . . S14
S5 Transcriptional promoter and terminator sequences . . . S14
S6 Plasmids used with terminator switch cgRNAs in *E. coli* . . . S16
S7 Plasmids used with splinted switch cgRNAs in *E. coli* . . . S19
S8 Plasmids used with toehold switch cgRNAs in *E. coli* . . . S22

### S1 Methods

#### S1.1 Rational design of libraries of orthogonal cgRNAs using NUPACK

For each mechanism, orthogonal cgRNA/trigger pairs were designed using the reaction pathway engineering tools within NUPACK (`nupack.org`; see the NUPACK 3.2 User Guide).[1,2] Target test tubes were specified using the general formulation of Section S2.2.1 in the Supplementary Information of Wolfe et al.[2] using the definitions provided below (Section S1.1.1 for the terminator switch, Section S1.1.2 for the splinted switch, and Section S1.1.3 for the toehold switch). Sequence designs were performed for libraries of 4 orthogonal cgRNA/trigger pairs. For a given design trial, the sequences were optimized by mutating the sequence set to reduce the multi-tube ensemble defect[2] subject to the diverse sequence constraints detailed below. Within the ensemble defect, defect weights (see Section S1.6 in the Supplementary Information of Wolfe et al.[2]) were applied to prioritize design effort as described below. Designs were performed using RNA parameters for 37 °C in 1M $Na^{+}$.[3] After performing several independent design trials for a given mechanism, a final sequence set was selected for experimental testing based on inspection of the predicted structural defects (fraction of nucleotides in the incorrect base-pairing state within the ensemble of an on-target complex) and concentration defects (fraction of nucleotides in the incorrect base-pairing state because there is a deficiency in the concentration of an on-target complex) for species in the context of the target test tubes,[2,4] as well as for each cgRNA in the presence of each non-cognate trigger (e.g., computational orthogonality study of Figure 5b (right)). After preliminary experimental studies, 3 cgRNA/trigger pairs (termed A, B, C) were selected for full experimental characterization.

##### S1.1.1 Target test tube specification for terminator switch mechanism

To design $N$ orthogonal systems, the total number of target test tubes is $|\Omega| = \sum_{n=1,...,N}$ {Step 0, Step 1}$_n$ + Crosstalk = $2N + 1$; the target test tubes in the multi-tube ensemble, $\Omega$, are indexed by $h = 1, \ldots, |\Omega|$. $L_{\max} = 2$ for all tubes (i.e., each target test tube contains all off-target complexes of up to 2 strands). Final sequence designs for orthogonal cgRNAs/triggers A, B, C are shown in Table S1.

###### Reactants for system *n*

- cgRNAs: $\text{G}_n$
- Triggers: $\text{X}_n$

###### Elementary step tubes for system *n*

- Step $0_n$ tube: $\Psi^{\text{products}}_{0_n} \equiv \{\text{G, X}\}_n$; $\Psi^{\text{reactants}}_{0_n} \equiv \emptyset$; $\Psi^{\text{exclude}}_{0_n} \equiv \{\text{G}{\cdot}\text{X}\}_n$
- Step $1_n$ tube: $\Psi^{\text{products}}_{1_n} \equiv \{\text{G}{\cdot}\text{X}\}_n$; $\Psi^{\text{reactants}}_{1_n} \equiv \{\text{G, X}\}_n$; $\Psi^{\text{exclude}}_{1_n} \equiv \emptyset$

###### Global crosstalk tube

- Crosstalk tube: $\Psi^{\text{reactive}}_{\text{global}} \equiv \cup_{n=1,...,N}\{\lambda^{\text{reactive}}_n\}$; $\Psi^{\text{crosstalk}}_{\text{global}} \equiv \Psi^{L \le L_{\max}}_{\text{global}} - \cup_{n=1,...,N}\{\lambda^{\text{cognate}}_n\}$

The reactive species and cognate products for system $n$ are:

- $\lambda^{\text{simple}}_n \equiv \{\text{G, X}\}_n$
- $\lambda^{\text{ss-out}}_n \equiv \text{X}_n$
- $\lambda^{\text{ss-in}}_n \equiv \text{G}^{\text{ss}}_n$, the 30nt single stranded terminator loop insert domain
- $\lambda^{\text{reactive}}_n \equiv \{\text{G, X, G}^{\text{ss}}\}_n$
- $\lambda^{\text{cognate}}_n \equiv \{\text{G}{\cdot}\text{X, G}^{\text{ss}}{\cdot}\text{X}\}_n$

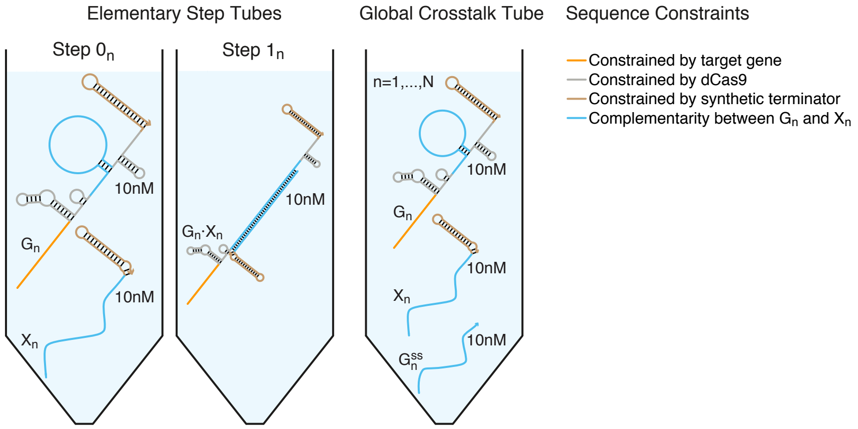


| Tube | On-targets ($\Psi^{\text{on}}_h$) | Off-targets ($\Psi^{\text{off}}_h$) |
|---|---|---|
| Step $0_n$ | $\{\text{G, X}\}_n$ | $\Psi^{L \le L_{\max}}_{0_n} - \{\text{G}{\cdot}\text{X}\}_n$ |
| Step $1_n$ | $\{\text{G}{\cdot}\text{X}\}_n$ | $\{\text{G, X}\}_n \cup \Psi^{L \le L_{\max}}_{1_n}$ |
| Crosstalk | $\cup_{n=1,...,N}\{\lambda^{\text{reactive}}_n\}$ | $\Psi^{L \le L_{\max}}_{\text{global}} - \cup_{n=1,...,N}\{\lambda^{\text{cognate}}_n\}$ |

**Figure S1: Target test tubes for sequence design of orthogonal terminator switch cgRNAs.** Top: Target test tube schematics. Bottom: Target test tube details. Each target test tube contains the depicted on-target complexes (each with the depicted target structure and a target concentration of 10 nM) and the off-target complexes listed in the table (each with vanishing target concentration). The on-target structures depicted above are used in the mechanism schematic of Figure 2a. To simultaneously design $N$ orthogonal systems, the total number of target test tubes is $|\Omega| = 2N + 1$. $L_{\max} = 2$ for all tubes. Domain shading reflects sequence constraints. Design conditions: RNA in 1 M $\text{Na}^+$ at 37 °C.

**Sequence constraints**

- Assignment constraints: portions of the cgRNA are constrained to match standard gRNA sequences for use with dCas9 (shaded gray in Figure 2a, Figure S1, and Table S1), the synthetic terminators for the cgRNA and trigger are fully constrained (shaded tan in Figure 2a, Figure S1, and Table S1).
- Watson–Crick constraints: cgRNA sequence domains "d-e-f" are constrained to be complementary to the trigger sequence domains "f*-e*-d*" (shaded blue in Figure 2a, Figure S1, and Table S1).
- Assignment constraint: cgRNA domain "u" is constrained to be complementary to a subsequence of the target gene mRFP (full template sequence in Section S2.3, constrained sequence shaded orange in Figure 2a, Figure S1, and Table S1).
- Pattern prevention constraints: the following patterns are prevented for cgRNA sequence domain "f": `AAAA`, `CCCC`, `GGGG`, `UUUU`.

**Defect weights**

- Test tube weight for each elementary step tube: 1
- Test tube weight for global crosstalk tube: 4
- Nucleotide weights are depicted for each complex in Figure S2

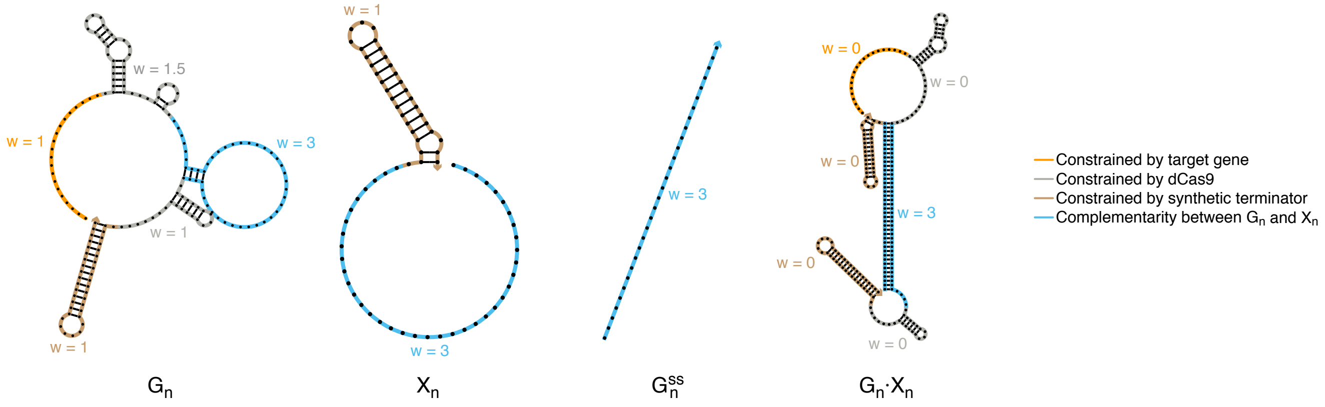


**Figure S2: Nucleotide defect weights for sequence design of terminator switch cgRNAs.** Within the target test tubes of Figure S1, the nucleotides in a given sequence domain within a given complex are assigned a defect weight *w* as depicted.

##### S1.1.2 Target test tube specification for splinted switch mechanism

To design $N$ orthogonal systems, the total number of target test tubes is $|\Omega| = \sum_{n=1,\ldots,N}$ {Step 0, Step 1}$_n$ + Crosstalk = $2N + 1$; the target test tubes in the multi-tube ensemble, $\Omega$, are indexed by $h = 1, \ldots, |\Omega|$. $L_{\max} = 2$ for all tubes (i.e., each target test tube contains all off-target complexes of up to 2 strands). Final sequence designs for orthogonal cgRNAs/triggers A, B, C are shown in Table S2.

###### Reactants for system *n*

- cgRNAs: $G_n$
- Triggers: $X_n$

###### Elementary step tubes for system *n*

- Step $0_n$ tube: $\Psi^{\text{products}}_{0_n} \equiv \{\text{G, X}\}_n$; $\Psi^{\text{reactants}}_{0_n} \equiv \emptyset$; $\Psi^{\text{exclude}}_{0_n} \equiv \{\text{G·X}\}_n$
- Step $1_n$ tube: $\Psi^{\text{products}}_{1_n} \equiv \{\text{G·X}\}_n$; $\Psi^{\text{reactants}}_{1_n} \equiv \{\text{G, X}\}_n$; $\Psi^{\text{exclude}}_{1_n} \equiv \emptyset$

###### Global crosstalk tube

- Crosstalk tube: $\Psi^{\text{reactive}}_{\text{global}} \equiv \cup_{n=1,\ldots,N}\{\lambda^{\text{reactive}}_n\}$; $\Psi^{\text{crosstalk}}_{\text{global}} \equiv \Psi^{L \le L_{\max}}_{\text{global}} - \cup_{n=1,\ldots,N}\{\lambda^{\text{cognate}}_n\}$

The reactive species and cognate products for system $n$ are:

- $\lambda^{\text{simple}}_n \equiv \{\text{G, X}\}_n$
- $\lambda^{\text{ss-out}}_n \equiv X_n$
- $\lambda^{\text{ss-in}}_n \equiv G^{\text{ss}}_n$, the 35nt single stranded handle and terminator loop insert domains with intervening gRNA sequence
- $\lambda^{\text{reactive}}_n \equiv \{\text{G, X, G}^{\text{ss}}\}_n$
- $\lambda^{\text{cognate}}_n \equiv \{\text{G·X, G}^{\text{ss}}\text{·X}\}_n$

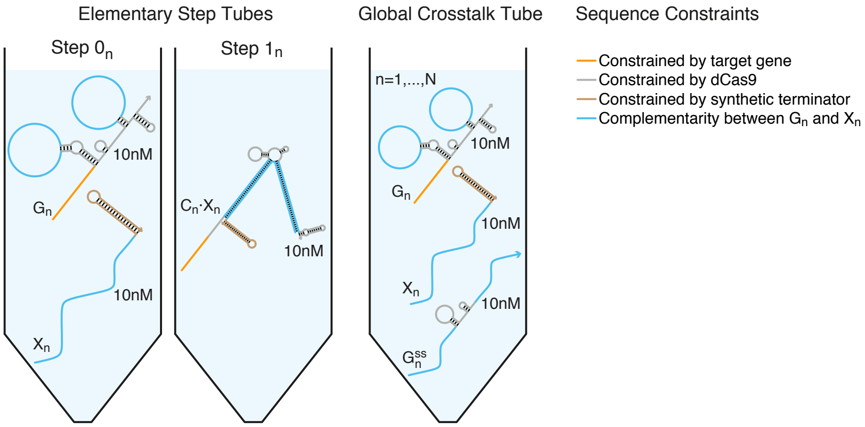


| Tube | On-targets ($\Psi^{\text{on}}_h$) | Off-targets ($\Psi^{\text{off}}_h$) |
|---|---|---|
| Step $0_n$ | $\{\text{G, X}\}_n$ | $\Psi^{L \le L_{\max}}_{0_n} - \{\text{G·X}\}_n$ |
| Step $1_n$ | $\{\text{G·X}\}_n$ | $\{\text{G, X}\}_n \cup \Psi^{L \le L_{\max}}_{1_n}$ |
| Crosstalk | $\cup_{n=1,\ldots,N}\{\lambda^{\text{reactive}}_n\}$ | $\Psi^{L \le L_{\max}}_{\text{global}} - \cup_{n=1,\ldots,N}\{\lambda^{\text{cognate}}_n\}$ |

**Figure S3: Target test tubes for sequence design of orthogonal splinted switch cgRNAs.** Top: Target test tube schematics. Bottom: Target test tube details. Each target test tube contains the depicted on-target complexes (each with the depicted target structure and a target concentration of 10 nM) and the off-target complexes listed in the table (each with vanishing target concentration). The on-target structures depicted above are used in the mechanism schematic of Figure 3a. To simultaneously design $N$ orthogonal systems, the total number of target test tubes is $|\Omega| = 2N + 1$. $L_{\max} = 2$ for all tubes. Design conditions: RNA in 1 M $Na^+$ at 37 °C.

##### Sequence constraints

- Assignment constraints: portions of the cgRNA are constrained to match standard gRNA sequences for use with dCas9 (shaded gray in Figure 3a, Figure S3, and Table S2), the synthetic terminator for the trigger is fully constrained (shaded tan in Figure 3a, Figure S3, and Table S2).
- Watson–Crick constraints: cgRNA sequence domains “d” and “e” are constrained to be complementary to the trigger sequence domains “d*” and “e*” (shaded blue in Figure 3a, Figure S3, and Table S2).
- Assignment constraint: cgRNA domain “u” is constrained to be complementary to a subsequence of the target gene sfGFP (full template sequence in Section S2.3, constrained sequence shaded orange in Figure 3a, Figure S3, and Table S2).
- Pattern prevention constraints: the following patterns are prevented for cgRNA sequence domains “d” and “e”: `AAAA`, `CCCC`, `GGGG`, `UUUU`.

##### Defect weights

- Test tube weight for each elementary step tube: 1
- Test tube weight for global crosstalk tube: 4
- Nucleotide weights are depicted for each complex in Figure S4

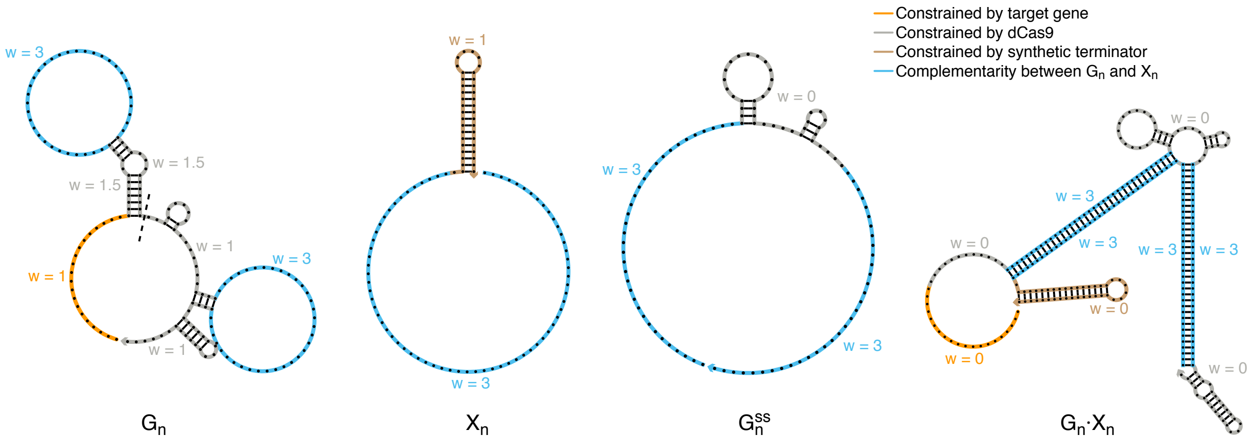


**Figure S4: Nucleotide defect weights for sequence design of splinted switch cgRNAs.** Within the target test tubes of Figure S3, the nucleotides in a given sequence domain within a given complex are assigned a defect weight *w* as depicted.

##### S1.1.3 Target test tube specification for toehold switch mechanism

Computational sequence design of the toehold switch mechanism was performed using a previous version of NUPACK that did not yet support exclusion of a set of complexes from a target test tube ensemble. For example, to design $N$ orthogonal systems, the test tube specifications detailed in the bottom tables of Figures S1 and S3 use the "minus" operator to enable compact specification of one Reactants tube for each system $n = 1, \ldots, N$ and a single Global Crosstalk tube. Lacking the minus operator at the time the toehold switch library was designed, we used a more verbose target test tube specification with separate Reactants tubes for cgRNA and trigger for each system $n = 1, \ldots, N$, as well as one Crosstalk tube for each non-cognate cgRNA/trigger pair.

To design $N$ orthogonal systems, the total number of target test tubes is:

$$|\Omega| = \sum_{n=1,\ldots,N} \{\text{Step } 0^{\text{G}}, \text{Step } 0^{\text{X}}, \text{Step } 1\}_n + \sum_{\substack{n=1,\ldots,N \\ p=1,\ldots,N \\ p\neq n}} \text{Crosstalk}_{p,n} \quad = 3N + N(N-1) = N^2 + 2N$$

The target test tubes in the multi-tube ensemble, $\Omega$, are indexed by $h = 1, \ldots, |\Omega|$. $L_{\max} = 2$ for all tubes (i.e., each target test tube contains all off-target complexes of up to 2 strands). Final sequence designs for orthogonal cgRNAs/triggers A, B, C are shown in Table S3.

**Reactants for system *n***

- cgRNAs: $\text{G}_n$
- Triggers: $\text{X}_n$

**Elementary step tubes for system *n***

- Step $0^{\text{G}}{}_n$ tube: $\Psi^{\text{products}}_{0_n} \equiv \text{G}_n$; $\Psi^{\text{reactants}}_{0_n} \equiv \emptyset$
- Step $0^{\text{X}}{}_n$ tube: $\Psi^{\text{products}}_{0_n} \equiv \text{X}_n$; $\Psi^{\text{reactants}}_{0_n} \equiv \emptyset$
- Step $1_n$ tube: $\Psi^{\text{products}}_{1_n} \equiv \{\text{G}\cdot\text{X}\}_n$; $\Psi^{\text{reactants}}_{1_n} \equiv \{\text{G, X}\}_n$

**Crosstalk tubes for system *n***

- Crosstalk tubes: $\Psi^{\text{products}}_{\text{crosstalk}_{p,n}} \equiv \{\text{G}_n, \text{X}_p\}$; $\Psi^{\text{reactants}}_{\text{crosstalk}_{p,n}} \equiv \emptyset$, for each non-cognate cgRNA/trigger pair ($p \neq n$)

**Sequence constraints**

- Assignment constraints: portions of the cgRNA are constrained to match standard gRNA sequences for use with dCas9 (shaded gray in Figure 4a, Figure S5 and Table S3), the synthetic terminator for the trigger is fully constrained (shaded tan in Figure 4a, Figure S5 and Table S3).
- Watson–Crick constraints: cgRNA sequence domains "d" is constrained to be complementary to the trigger sequence domains "d*" (shaded blue in Figure 4a, Figure S5 and Table S3).
- Assignment constraint: cgRNA domain "u" is constrained to be complementary to a subsequence of the target gene mRFP (full template sequence in Section S2.3, constrained sequence shaded orange in Figure 4a, Figure S5 and Table S3).
- Pattern prevention constraints: the following patterns are prevented for cgRNA sequence domain "d": `AAAA, CCCC, GGGG, UUUU, KKKKKK, MMMMMM, RRRRRR, SSSSSS, WWWWWW, YYYYYY.`

**Defect weights**

- Test tube weight for each elementary step tube: 1
- Test tube weight for crosstalk tubes: 1
- Nucleotide weights are depicted for each complex in Figure S6

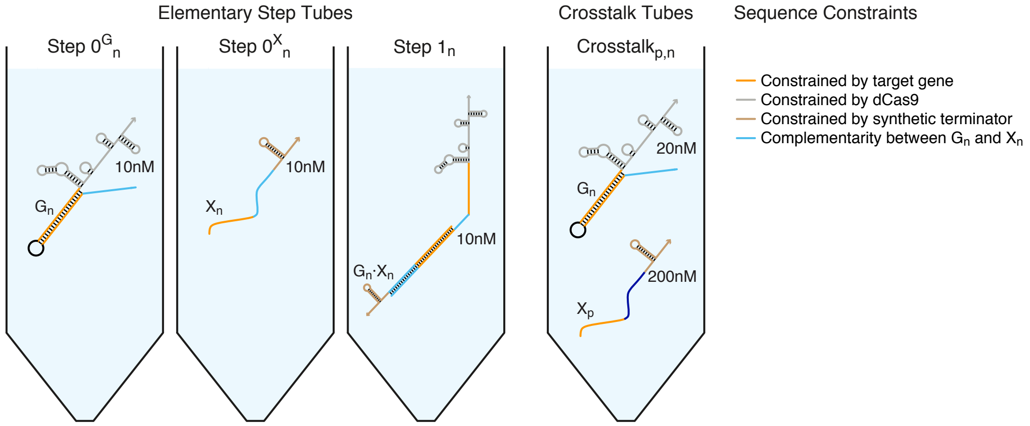


| Tube | On-targets ($\Psi_h^{\text{on}}$) | Off-targets ($\Psi_h^{\text{off}}$) |
|---|---|---|
| Step $0^{\text{G}}_n$ | $\text{G}_n$ | $\Psi^{L\le L_{\max}}_{0^G_n}$ |
| Step $0^{\text{X}}_n$ | $\text{X}_n$ | $\Psi^{L\le L_{\max}}_{0^X_n}$ |
| Step $1_n$ | $\{\text{G}\cdot\text{X}\}_n$ | $\{\text{G},\text{X}\}_n \cup \Psi^{L\le L_{\max}}_{1_n}$ |
| Crosstalk$_{p,n}$ | $\{\text{G}_n, \text{X}_p\}$ | $\{\text{G}_n\cdot\text{X}_p\} \cup \Psi^{L\le L_{\max}}_{\text{crosstalk}_{p,n}}$ |

**Figure S5: Target test tubes for sequence design of orthogonal toehold switch cgRNAs.** Top: Target test tube schematics. Bottom: Target test tube details. Each target test tube contains the depicted on-target complexes (each with the depicted target structure and target concentration) and the off-target complexes listed in the table (each with vanishing target concentration). The on-target structures depicted above are used in the mechanism schematic of Figure 4a. To simultaneously design $N$ orthogonal systems, the total number of target test tubes is $|\Omega| = N^2 + 2N$. $L_{\max} = 2$ for all tubes. Design conditions: RNA in 1 M $\text{Na}^+$ at 37 °C.

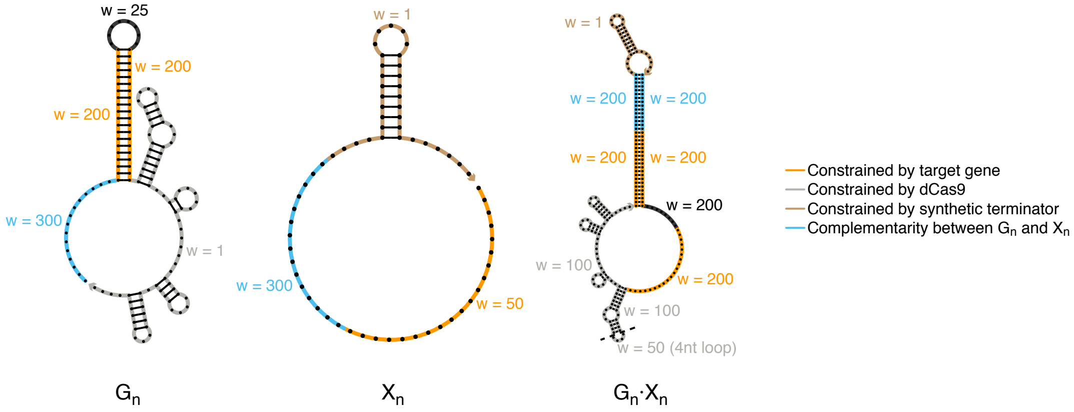


**Figure S6: Nucleotide defect weights for sequence design of toehold switch cgRNAs.** Within the target test tubes of Figure S5, the nucleotides in a given sequence domain within a given complex are assigned a defect weight *w* as depicted.

#### S1.2 Plasmid construction and molecular cloning

Sequences for the parts used in this study are provided in Section S2. Plasmid layouts for each construct as well as example plasmid maps and corresponding full sequences are provided in Section S3.

Control gRNA and cgRNA constructs were generated by inverse PCR, inserting sequence modifications into the previously described pgRNA-bacteria vector[5] (Addgene plasmid #44251; gift from Dr. Stanley Qi). All PCR steps for the generation of experimental constructs were performed using Q5 Hot Start High-Fidelity polymerase (NEB #M0494) according to manufacturer instructions using primers designed using standard molecular cloning techniques and synthesized by Integrated DNA Technologies. Introduced sequences were verified by Sanger sequencing for single colony picks via colony PCR using GoTaq Green polymerase (Promega #M7122).

For terminator switch cgRNAs (Figure 2), trigger-expressing constructs were generated by cloning lacI-regulated promoter (BioBrick part number BBa_R0011), trigger template, and synthetic terminator (BBa_B1002) directly into the cgRNA vector via inverse PCR. For splinted switch cgRNAs (Figure 3), trigger-expressing constructs were generated by first cloning synthetic promoter, trigger template, and synthetic terminator (BBa_B1006) into a trigger-only cassette via inverse PCR, and then cgRNA+trigger expressing constructs were cloned by inserting trigger cassette into the cgRNA vector using BioBrick assembly.[6,7] For toehold switch cgRNAs (Figure 4), trigger-expressing constructs were generated by first cloning synthetic promoter, trigger template, and synthetic terminator (BBa_B0050) into a trigger-only cassette via inverse PCR, and then cgRNA+trigger expressing constructs were cloned by inserting trigger cassette into the cgRNA vector using DNA assembly according to manufacturer instructions (NEBuilder HiFi DNA Assembly, NEB #E2621).

A lacI+dCas9 expression construct was generated by inserting a lacI template sequence with J23108 constitutive promoter[8] into the previously described pdCas9-bateria vector[5] (Addgene plasmid #44249; gift from Dr. Stanley Qi) between the dCas9 gene and the p15A origin with a synthetic terminator (BBa_B0010) added upstream of lacI as a transcriptional terminator for dCas9 (see Section S3.4), using DNA assembly according to manufacturer instructions (NEBuilder HiFi DNA Assembly, NEB #E2621).

#### S1.3 Bacterial culture for cgRNA studies

A previously described *E. coli* MG1655 strain with constitutively expressed mRFP and sfGFP inserted into the nfsA locus[5] (Ec001; gift from Dr. Stanley Qi) was used for all fluorescence assays. For experiments with constitutive expression of trigger, the previously described pdCas9-bacteria vector[5] (Addgene plasmid #44251) was used for tetR-regulated dCas9 expression. For experiments with lacI-regulated expression of trigger, the lacI+dCas9 vector was used for tetR-regulated dCas9 expression and constitutive expression of lacI. Chemically competent chloramphenicol-resistant cells carrying either the dCas9 or lacI+dCas9 construct were transformed with gRNA, cgRNA, or cgRNA+trigger expression vectors and cultivated in EZ-RDM (Teknova #M2105) containing 100 $\mu$g/mL carbenicillin and 34 $\mu$g/mL chloramphenicol (EZ-RDM+Carb+Cam).

Sequence-verified strains were grown overnight in EZ-RDM+Carb+Cam, then seeded at 100$\times$ dilution in 100 $\mu$L fresh medium and grown at 37 °C with shaking in the Neo2 microplate reader (Biotek) to monitor absorbance at 600 nm. When cells had reached mid-log phase ($\approx$4 h), cells were again diluted $\approx$100-fold in fresh medium with cell density normalized by A600 and, if applicable, dCas9 expression and trigger expression were induced with aTc and IPTG, respectively, in $N = 3$ replicate wells at 400 $\mu$L final volume in a 96-well high-volume glass bottom plate (Matriplate, Brooks #MGB096-1-2-LG-L). A final working concentration of 200 nM aTc and 5mM IPTG was used for terminator switch experiments (Figure 2). A final working concentration of 2 nM aTc was used for splinted switch experiments (Figure 3). A final working concentration of 200 nM aTc was used for toehold switch experiments (Figure 4). Induced cells were grown at 37 °C with continuous shaking for 12 h.

#### S1.4 Flow cytometry

Protein fluorescence was measured using the MACSQuant VYB flow cytometer (Miltenyi Biotec) using FSC/SSC to gate for 20,000 live cells per well (see example of Figure S7) at a flow rate of 25 $\mu$L/min. sfGFP fluorescence was measured using the B1 channel (488 nm laser, 525/50 nm filter) and mRFP fluorescence was measured using the Y2 channel (561 nm laser, 615/20 nm filter).

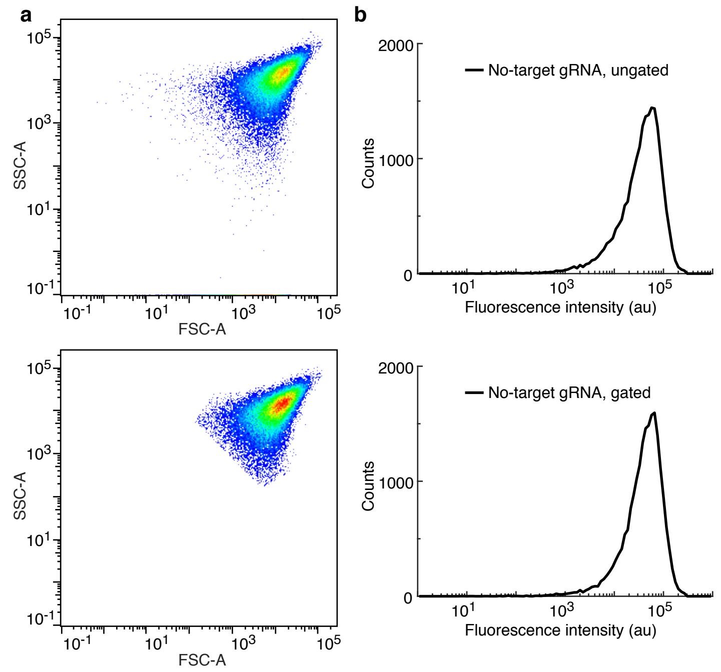


**Figure S7: Illustration of gates used for flow cytometry analysis of *E. coli.*** (a) Scatter plots for ungated sample (top) and gated sample (bottom): side scatter area (SSC-A) vs forward scatter area (FSC-A). (b) Fluorescence intensity histogram for ungated sample (top) and gated sample (bottom). mRFP fluorescence for no-target gRNA control used for terminator switch characterization in Figure 2b.

#### S1.5 Cell fluorescence analysis

For a bacterial strain containing a fluorescent reporter protein, the total fluorescence (SIG+AF) in the relevant fluorescent channel (mRFP for the terminator switch of Figure 2, sfGFP for the splinted switch of Figure 3, mRFP for the toehold switch of Figure 4) is a combination of signal (SIG) from the reporter and autofluorescence (AF) inherent to the cells. Autofluorescence is characterized for a given fluorescent channel in strain MG1655 containing no fluorescent reporters. For cell $j$ of replicate well $i$ of a given bacterial strain, we denote the autofluorescence:

$$X_{i,j}^{\mathrm{AF}}$$

the signal

$$X_{i,j}^{\mathrm{SIG}}$$

and the total fluorescence (SIG + AF):

$$X_{i,j}^{\mathrm{SIG+AF}}$$

For replicate well $i$ of a given strain, we measure the median fluorescence ($\tilde{X}_i^{\mathrm{SIG+AF}}$ for a strain containing the reporter, $\tilde{X}_i^{\mathrm{AF}}$ for strain MG1655 lacking reporters) over $N = 20,000$ cells. Performance across $N = 3$ replicate wells is characterized by the sample means ($\bar{X}^{\mathrm{SIG+AF}}$ and $\bar{X}^{\mathrm{AF}}$) and standard errors ($s_{\bar{X}^{\mathrm{SIG+AF}}}$ and $s_{\bar{X}^{\mathrm{AF}}}$). Let $(n, p)$ denote a strain containing cgRNA $n$ and trigger $p$. The mean signal is estimated as

$$\bar{X}(n,p)^{\mathrm{SIG}} = \bar{X}(n,p)^{\mathrm{SIG+AF}} - \bar{X}^{\mathrm{AF}}$$

with the standard error estimated via uncertainty propagation as

$$s_{\bar{X}(n,p)^{\mathrm{SIG}}} \leq \sqrt{(s_{\bar{X}(n,p)^{\mathrm{SIG+AF}}})^2 + (s_{\bar{X}^{\mathrm{AF}}})^2}.$$

The upper bound on estimated standard error holds under the assumption that the correlation between SIG and AF is non-negative.

###### S1.5.1 OFF:ON ratio for constitutively active cgRNAs (ON→OFF logic)

For a constitutively active cgRNA with silencing dCas9 (Figures 2 and 3), the ON state for cgRNA $n$ corresponds to low fluorescence using no trigger ($p = 0$) and the OFF state corresponds to high fluorescence using cognate trigger $p = n$. The OFF:ON ratio is estimated as

$$\bar{X}(n)^{\text{OFF:ON}} = \bar{X}(n,n)^{\text{SIG}} / \bar{X}(n,0)^{\text{SIG}}$$

with standard error estimated via uncertainty propagation as

$$s(n)^{\text{OFF:ON}} \leq \bar{X}(n)^{\text{OFF:ON}} \sqrt{\left(\frac{s_{\bar{X}(n,n)^{\text{SIG}}}}{\bar{X}(n,n)^{\text{SIG}}}\right)^2 + \left(\frac{s_{\bar{X}(n,0)^{\text{SIG}}}}{\bar{X}(n,0)^{\text{SIG}}}\right)^2}$$

The upper bound on estimated standard error holds under the assumption that the correlation between SIG in the two strains is non-negative.

###### S1.5.2 OFF:ON ratio for constitutively inactive cgRNAs (OFF→ON logic)

For a constitutively inactive cgRNA with silencing dCas9 (Figure 4), the OFF state for cgRNA $n$ corresponds to high fluorescence using no trigger $p = 0$ and the ON state corresponds to low fluorescence using cognate trigger ($p = n$). The OFF:ON ratio is estimated as

$$\bar{X}(n)^{\text{OFF:ON}} = \bar{X}(n,0)^{\text{SIG}} / \bar{X}(n,n)^{\text{SIG}}$$

with standard error estimated via uncertainty propagation as

$$s(n)^{\text{OFF:ON}} \leq \bar{X}(n)^{\text{OFF:ON}} \sqrt{\left(\frac{s_{\bar{X}(n,0)^{\text{SIG}}}}{\bar{X}(n,0)^{\text{SIG}}}\right)^2 + \left(\frac{s_{\bar{X}(n,n)^{\text{SIG}}}}{\bar{X}(n,n)^{\text{SIG}}}\right)^2}$$

The upper bound on estimated standard error holds under the assumption that the correlation between SIG in the two strains is non-negative. In the bar graphs of Figures 2c (left), 3c (left), 4c (left), we report mean $\pm$ estimated standard error of the mean of median single-cell fluorescence over 20,000 cells for $N = 3$ replicate wells.

###### S1.5.3 Crosstalk for orthogonal cgRNAs

Crosstalk (CT) is estimated for cgRNA $n$ with trigger $p$ as

$$\bar{X}(n,p)^{\text{CT}} = [\bar{X}(n,p)^{\text{SIG}} - \bar{X}(n,0)^{\text{SIG}}] / [\bar{X}(n,n)^{\text{SIG}} - \bar{X}(n,0)^{\text{SIG}}].$$

with the standard error estimated via uncertainty propagation as

$$s(n,p)^{\text{CT}} \leq \bar{X}(n,p)^{\text{CT}} \sqrt{\left(\frac{\sqrt{(s_{\bar{X}(n,p)^{\text{SIG}}})^2 + (s_{\bar{X}(n,0)^{\text{SIG}}})^2}}{\bar{X}(n,p)^{\text{SIG}} - \bar{X}(n,0)^{\text{SIG}}}\right)^2 + \left(\frac{\sqrt{(s_{\bar{X}(n,n)^{\text{SIG}}})^2 + (s_{\bar{X}(n,0)^{\text{SIG}}})^2}}{\bar{X}(n,n)^{\text{SIG}} - \bar{X}(n,0)^{\text{SIG}}}\right)^2}$$

The upper bound on estimated standard error holds under the assumption that the correlation between SIG in the two strains is non-negative. Note that crosstalk values can be positive or negative. In the bar graphs of Figures 2c (right), 3c (right), 4c (right), we report mean $\pm$ estimated standard error of the mean ($\bar{X} \pm s$) of median single-cell fluorescence over 20,000 cells for $N = 3$ replicate wells. In instances where $|\bar{X}| < s$, we instead report ($\bar{X} + s$) as an estimated upper bound.

### S2 Sequences

#### S2.1 Sequences for cgRNAs, triggers, and control gRNAs

| Name | Sequence | Figure | Legend |
|---|---|---|---|
| TrmS_cgA | 5′-AACTTTCAGTTTAGCGGTCTGTTTTAGAGCTAGAAATAGCAAGTTAAAAT<br>AAGGCTAGTCCGATCAAACGGGTAAACAAACAGGATAATTAAGGAGGCAGTA<br>CCCGGGCACCGAGTCGGTGCTTTTTTT-3′ | 2b<br>2c | cgRNA<br>cgRNA A |
| TrmS_cgB | 5′-AACTTTCAGTTTAGCGGTCTGTTTTAGAGCTAGAAATAGCAAGTTAAAAT<br>AAGGCTAGTCCGTATCATGGGGTTGTGTGTTGTTGTAAGTGTGTGTGTGTTG<br>CCCCGGCACCGAGTCGGTGCTTTTTTT-3′ | 2c | cgRNA B |
| TrmS_cgC | 5′-AACTTTCAGTTTAGCGGTCTGTTTTAGAGCTAGAAATAGCAAGTTAAAAT<br>AAGGCTAGTCCGAATATAGGGGAAGAGAAAGAAGAAGAGAAGAGAAAGATGT<br>CCCCGGCACCGAGTCGGTGCTTTTTTT-3′ | 2c | cgRNA C |
| TrmS_tA | 5′-TACTGCCTCCTTAATTATCCTGTTTGTTTACCCGTTTGAT-3′ | 2b<br>2c | Trigger<br>Trigger A |
| TrmS_tB | 5′-CAACACACACACACTTACAACAACACACAACCCCATGATA-3′ | 2c | Trigger B |
| TrmS_tC | 5′-ACATCTTTCTCTTCTCTTCTTCTTTCTCTTCCCCTATATT-3′ | 2c | Trigger C |

**Table S1: Terminator switch sequences.** Nucleotides shaded orange are constrained by the target gene. Nucleotides shaded gray are constrained by dCas9. Nucleotides shaded blue are designed as described in Section S1.1.

| Name | Sequence | Figure | Legend |
|---|---|---|---|
| SplS_cgA | 5′-CATCTAATTCAACAAGAATTGTTTTAGAGCTACACCTTACGCCGGTTCAA<br>TTCCAAGTCCCTTCCAGTAGCAAGTTAAAATAAGGCTAGTCCGTTATCAACT<br>TAACACCCTTTACAAACCTTCCTCTTCCTTTACCCTAAGTGGCACCGAGTCG<br>GTGCTTTTTTT-3′ | 3b<br>3c | cgRNA<br>cgRNA A |
| SplS_cgB | 5′-CATCTAATTCAACAAGAATTGTTTTAGAGCTAGTAATCGAATCATAGTAA<br>ATTTCCCATCGTCATAATAGCAAGTTAAAATAAGGCTAGTCCGTTATCAACT<br>TCATACGGGTCTGAAGTAGTTCATTCTTATACAGTCAAGTGGCACCGAGTCG<br>GTGCTTTTTTT-3′ | 3c | cgRNA B |
| SplS_cgC | 5′-CATCTAATTCAACAAGAATTGTTTTAGAGCTAGTCGTTACCTTATCAATA<br>TCAACCTCCGCATACACTAGCAAGTTAAAATAAGGCTAGTCCGTTATCAACT<br>TGCACATAGGACCCAACATGCCAACAGAGAAGAGTTAAGTGGCACCGAGTCG<br>GTGCTTTTTTT-3′ | 3c | cgRNA C |
| SplS_tA | 5′-AGGGTAAAGGAAGAGGAAGGTTTGTAAAGGGTGTTCTGGAAGGGACTTGG<br>AATTGAACCGGCGTAAGGTG-3′ | 3b<br>3c | Trigger<br>Trigger A |
| SplS_tB | 5′-GACTGTATAAGAATGAACTACTTCAGACCCGTATGTTATGACGATGGGAA<br>ATTTACTATGATTCGATTAC-3′ | 3c | Trigger B |
| SplS_tC | 5′-AACTCTTCTCTGTTGGCATGTTGGGTCCTATGTGCGTGTATGCGGAGGTT<br>GATATTGATAAGGTAACGAC-3′ | 3c | Trigger C |

**Table S2: Splinted switch sequences.** Nucleotides shaded orange are constrained by the target gene. Nucleotides shaded gray are constrained by dCas9. Nucleotides shaded blue are designed as described in Section S1.1.

| Name | Sequence | Figure | Legend |
|---|---|---|---|
| ToeS_cgA | 5′−ATGTTCGTTGTATTAAGACCGCTAAACTGAAAGTTACACGCCCAACTTTC<br>AGTTTAGCGGTCTGTTTTAGAGCTAGAAATAGCAAGTTAAAATAAGGCTAGT<br>CCGTTATCAACTTGAAAAAGTGGCACCGAGTCGGTGCTTTTTTT−3′ | 4b<br>4c | cgRNA<br>cgRNA A |
| ToeS_cgB | 5′−GTATATGAAATTGAAAGACCGCTAAACTGAAAGTTACACGCCCAACTTTC<br>AGTTTAGCGGTCTGTTTTAGAGCTAGAAATAGCAAGTTAAAATAAGGCTAGT<br>CCGTTATCAACTTGAAAAAGTGGCACCGAGTCGGTGCTTTTTTT−3′ | 4c | cgRNA B |
| ToeS_cgC | 5′−AAGGTGATAGTAAAGAGACCGCTAAACTGAAAGTTACACGCCCAACTTTC<br>AGTTTAGCGGTCTGTTTTAGAGCTAGAAATAGCAAGTTAAAATAAGGCTAGT<br>CCGTTATCAACTTGAAAAAGTGGCACCGAGTCGGTGCTTTTTTT−3′ | 4c | cgRNA C |
| ToeS_tA | 5′−AACTTTCAGTTTAGCGGTCTTAATACAACGAACAT−3′ | 4b<br>4c | Trigger<br>Trigger A |
| ToeS_tB | 5′−AACTTTCAGTTTAGCGGTCTTTCAATTTCATATAC−3′ | 4c | Trigger B |
| ToeS_tC | 5′−AACTTTCAGTTTAGCGGTCTCTTTACTATCACCTT−3′ | 4c | Trigger C |

**Table S3: Toehold switch sequences.** Nucleotides shaded orange are constrained by the target gene. Nucleotides shaded gray are constrained by dCas9. Nucleotides shaded blue are designed as described in Section S1.1.

| Name | Sequence | Figure | Legend |
|---|---|---|---|
| mRFP_g | 5′−AACTTTCAGTTTAGCGGTCTGTTTTAGAGCTAGAAATAGCAAGTTAAAAT<br>AAGGCTAGTCCGTTATCAACTTGAAAAAGTGGCACCGAGTCGGTGCTTTTTT<br>T−3′ | 2b<br>4b | Standard gRNA<br>Standard gRNA |
| sfGFP_g | 5′−CATCTAATTCAACAAGAATTGTTTTAGAGCTAGAAATAGCAAGTTAAAAT<br>AAGGCTAGTCCGTTATCAACTTGAAAAAGTGGCACCGAGTCGGTGCTTTTTT<br>T−3′ | 3b | Standard gRNA |
| NT_g | 5′−GTTTTAGAGCTAGAAATAGCAAGTTAAAATAAGGCTAGTCCGTTATCAAC<br>TTGAAAAAGTGGCACCGAGTCGGTGCTTTTTTT−3′ | 2b<br>3b<br>4b | No-target gRNA, Autofluorescence<br>No-target gRNA, Autofluorescence<br>No-target gRNA, Autofluorescence |

**Table S4: Control gRNA sequences.** For the standard gRNAs, nucleotides shaded orange are constrained by the target gene (mRFP or sfGFP). The no-target gRNA contains no target-binding region. The autofluorescence control strain was transformed with the no-target gRNA control plasmid (sequence NT_g).

#### S2.2 Transcriptional promoter and terminator sequences

| Name | Type | Sequence |
|---|---|---|
| BBa_J23100 | Constitutive Promoter | 5′−TTGACGGCTAGCTCAGTCCTAGGTACAGTGCTAGC−3′ |
| BBa_J23108 | Constitutive Promoter | 5′−CTGACAGCTAGCTCAGTCCTAGGTATAATGCTAGC−3′ |
| BBa_J23114 | Constitutive Promoter | 5′−TTTATGGCTAGCTCAGTCCTAGGTACAATGCTAGC−3′ |
| BBa_R0011 | lacI Promoter | 5′−AATTGTGAGCGGATAACAATTGACATTGTGAGCGGATAACAAGATACTGAGCACA−3′ |
| BBa_B0015 | Synthetic Terminator | 5′−CCAGGCATCAAATAAAACGAAAGGCTCAGTCGAAAGACTGGGCCTTTCGTTTTATCT<br>GTTGTTTGTCGGTGAACGCTCTCTACTAGAGTCACACTGGCTCACCTTCGGGTGGGCCT<br>TTCTGCGTTTATA−3′ |
| BBa_B0050 | Synthetic Terminator | 5′−AAAAAAAGGATCTCAAGAAGATCCTTTGATTTT−3′ |
| BBa_B1002 | Synthetic Terminator | 5′−CGCAAAAAACCCCGCTTCGGCGGGGTTTTTTCGC−3′ |
| BBa_B1006 | Synthetic Terminator | 5′−AAAAAAAAACCCCGCCCCTGACAGGGCGGGGTTTTTTTT−3′ |
| BBa_B1010 | Synthetic Terminator | 5′−CGCCGCAAACCCCGCCCCTGACAGGGCGGGGTTTCGCCGC−3′ |

**Table S5: Transcriptional promoter and terminator sequences.**

#### S2.3 Gene sequences

##### mRFP template sequence

Nucleotides highlighted in orange indicate the gRNA/cgRNA target sequence.

```
1 ATGGCGAGTA GCGAAGACGT TATCAAAGAG TTCATGCGTT TCAAAGTTCG TATGGAAGGT TCCGTTAACG
```

71 GTCACGAGTT CGAAATCGAA GGTGAAGGTG AAGGTCGTCC GTACGAAGGT ACCCAGACCG CTAAACTGAA
141 AGTTACCAAA GGTGGTCCGC TGCCGTTCGC TTGGGACATC CTGTCCCCGC AGTTCCAGTA CGGTTCCAAA
211 GCTTACGTTA AACACCCGGC TGACATCCCG GACTACCTGA AACTGTCCTT CCCGGAAGGT TTCAAATGGG
281 AACGTGTTAT GAACTTCGAA GACGGTGGTG TTGTTACCGT TACCCAGGAC TCCTCCCTGC AAGACGGTGA
351 GTTCATCTAC AAAGTTAAAC TGCGTGGTAC CAACTTCCCG TCCGACGGTC CGGTTATGCA GAAAAAAACC
421 ATGGGTTGGG AAGCTTCCAC CGAACGTATG TACCCGGAAG ACGGTGCTCT GAAAGGTGAA ATCAAAATGC
491 GTCTGAAACT GAAAGACGGT GGTCACTACG ACGCTGAAGT TAAAACCACC TACATGGCTA AAAAACCGGT
561 TCAGCTGCCG GGTGCTTACA AAACCGACAT CAAACTGGAC ATCACCTCCC ACAACGAAGA CTACACCATC
631 GTTGAACAGT ACGAACGTGC TGAAGGTCGT CACTCCACCG GTGCTTAA

**sfGFP template sequence**

Nucleotides highlighted in orange indicate the gRNA/cgRNA target sequence.

1 ATGAGCAAAG GAGAAGAACT TTTCACTGGA GTTGTCCCAA TTCTTGTTGA ATTAGATGGT GATGTTAATG
71 GGCACAAATT TTCTGTCCGT GGAGAGGGTG AAGGTGATGC TACAAACGGA AAACTCACCC TTAAATTTAT
141 TTGCACTACT GGAAAACTAC CTGTTCCGTG GCCAACACTT GTCACTACTC TGACCTATGG TGTTCAATGC
211 TTTTCCCGTT ATCCGGATCA CATGAAACGG CATGACTTTT TCAAGAGTGC CATGCCCGAA GGTTATGTAC
281 AGGAACGCAC TATATCTTTC AAAGATGACG GGACCTACAA GACGCGTGCT GAAGTCAAGT TTGAAGGTGA
351 TACCCTTGTT AATCGTATCG AGTTAAAGGG TATTGATTTT AAAGAAGATG GAAACATTCT TGGACACAAA
421 CTCGAGTACA ACTTTAACTC ACACAATGTA TACATCACGG CAGACAAACA AAGAATGGA ATCAAAGCTA
491 ACTTCAAAAT TCGCCACAAC GTTGAAGATG GTTCCGTTCA ACTAGCAGAC CATTATCAAC AAAATACTCC
561 AATTGGCGAT GGCCCTGTCC TTTTACCAGA CAACCATTAC CTGTCGACAC AATCTGTCCT TTCGAAAGAT
631 CCCAACGAAA AGCGTGACCA CATGGTCCTT CTTGAGTTTG TAACTGCTGC TGGGATTACA CATGGCATGG
701 ATGAGCTCTA CAAA

**lacI template sequence[8]**

1 ATGGTGAATG TGAAACCAGT AACGTTATAC GATGTCGCAG AGTATGCCGG TGTCTCTTAT CAGACCGTTT
71 CCCGCGTGGT GAACCAGGCC AGCCACGTTT CTGCGAAAAC GCGGGAAAAA GTGGAAGCGG CGATGGCGGA
141 GCTGAATTAC ATTCCCAACC GCGTGGCACA ACAACTGGCG GGCAAACAGT CGTTGCTGAT TGGCGTTGCC
211 ACCTCCAGTC TGGCCCTGCA CGCGCCGTCG CAAATTGTCG CGGCGATTAA ATCTCGCGCC GATCAACTGG
281 GTGCCAGCGT GGTGGTGTCG ATGGTAGAAC GAAGCGGCGT CGAAGCCTGT AAAGCGGCGG TGCACAATCT
351 TCTCGCGCAA CGCGTCAGTG GGCTGATCAT TAACTATCCG CTGGATGACC AGGATGCCAT TGCTGTGGAA
421 GCTGCCTGCA CTAATGTTCC GGCGTTATTT CTTGATGTCT CTGACCAGAC ACCCATCAAC AGTATTATTT
491 TCTCCCATGA GGACGGTACG CGACTGGGCG TGGAGCATCT GGTCGCATTG GGTCACCAGC AAATCGCGCT
561 GTTAGCGGGC CCATTAAGTT CTGTCTCGGC GCGTCTGCGT CTGGCTGGCT GGCATAAATA TCTCACTCGC
631 AATCAAATTC AGCCGATAGC GGAACGGGAA GGCGACTGGA GTGCCATGTC CGGTTTTCAA CAAACCATGC
701 AAATGCTGAA TGAGGGCATC GTTCCCACTG CGATGCTGGT TGCCAACGAT CAGATGGCGC TGGGCGCAAT
771 GCGCGCCATT ACCGAGTCCG GGCTGCGCGT TGGTGCGGAT ATCTCGGTAG TGGGATACGA CGATACCGAA
841 GATAGCTCAT GTTATATCCC GCCGTTAACC ACCATCAAAC AGGATTTTCG CCTGCTGGGG CAAACCAGCG
911 TGGACCGCTT GCTGCAACTC TCTCAGGGCC AGGCGGTGAA GGGCAATCAG CTGTTGCCCG TCTCACTGGT
981 GAAAAGAAAA ACCACCCTGG CGCCCAATAC GCAAACCGCC TCTCCCCGCG CGTTGGCCGA TTCATTAATG
1051 CAGCTGGCAC GACAGGTTTC CCGACTGGAA AGCGGGCAGT AA

### S3 Plasmids

#### S3.1 Constitutively active terminator switch in *E. coli*

| Name | Parts | Figure | Legend |
|---|---|---|---|
| pdCas9+lacI | See Figure S14, Figure S15 | 2bc | |
| pg-NT | ColE1 ori; amp-R; BBa_J23114-NT_g-BBa_B1006 | 2b | No-target gRNA; Autofluorescence |
| | | 2c | Autofluorescence |
| pg-mRFP | ColE1 ori; amp-R; BBa_J23114-mRFP_g-BBa_B1006 | 2b | Standard gRNA |
| pTrmS-cgA-nT | ColE1 ori; amp-R; BBa_J23114-TrmS_cgA-BBa_B1006; BBa_R0011-BBa_B1002 | 2b | cgRNA |
| | | 2c | cgRNA A |
| pTrmS-cgA-tA | ColE1 ori; amp-R; BBa_J23114-TrmS_cgA-BBa_B1006; BBa_R0011-TrmS_tA-BBa_B1002 | 2b | cgRNA + trigger |
| | See Figure S8, Figure S9 | 2c | cgRNA A, trigger $X_A$ |
| pTrmS-cgA-tB | ColE1 ori; amp-R; BBa_J23114-TrmS_cgA-BBa_B1006; BBa_R0011-TrmS_tB-BBa_B1002 | 2c | cgRNA A, trigger $X_B$ |
| pTrmS-cgA-tC | ColE1 ori; amp-R; BBa_J23114-TrmS_cgA-BBa_B1006; BBa_R0011-TrmS_tC-BBa_B1002 | 2c | cgRNA A, trigger $X_C$ |
| pTrmS-cgB-nT | ColE1 ori; amp-R; BBa_J23114-TrmS_cgB-BBa_B1006; BBa_R0011-BBa_B1002 | 2c | cgRNA B |
| pTrmS-cgB-tA | ColE1 ori; amp-R; BBa_J23114-TrmS_cgB-BBa_B1006; BBa_R0011-TrmS_tA-BBa_B1002 | 2c | cgRNA B, trigger $X_A$ |
| pTrmS-cgB-tB | ColE1 ori; amp-R; BBa_J23114-TrmS_cgB-BBa_B1006; BBa_R0011-TrmS_tB-BBa_B1002 | 2c | cgRNA B, trigger $X_B$ |
| pTrmS-cgB-tC | ColE1 ori; amp-R; BBa_J23114-TrmS_cgB-BBa_B1006; BBa_R0011-TrmS_tC-BBa_B1002 | 2c | cgRNA B, trigger $X_C$ |
| pTrmS-cgC-nT | ColE1 ori; amp-R; BBa_J23114-TrmS_cgC-BBa_B1006; BBa_R0011-BBa_B1002 | 2c | cgRNA C |
| pTrmS-cgC-tA | ColE1 ori; amp-R; BBa_J23114-TrmS_cgC-BBa_B1006; BBa_R0011-TrmS_tA-BBa_B1002 | 2c | cgRNA C, trigger $X_A$ |
| pTrmS-cgC-tB | ColE1 ori; amp-R; BBa_J23114-TrmS_cgC-BBa_B1006; BBa_R0011-TrmS_tB-BBa_B1002 | 2c | cgRNA C, trigger $X_B$ |
| pTrmS-cgC-tC | ColE1 ori; amp-R; BBa_J23114-TrmS_cgC-BBa_B1006; BBa_R0011-TrmS_tC-BBa_B1002 | 2c | cgRNA C, trigger $X_C$ |

**Table S6: Plasmids used with terminator switch cgRNAs in *E. coli*.**

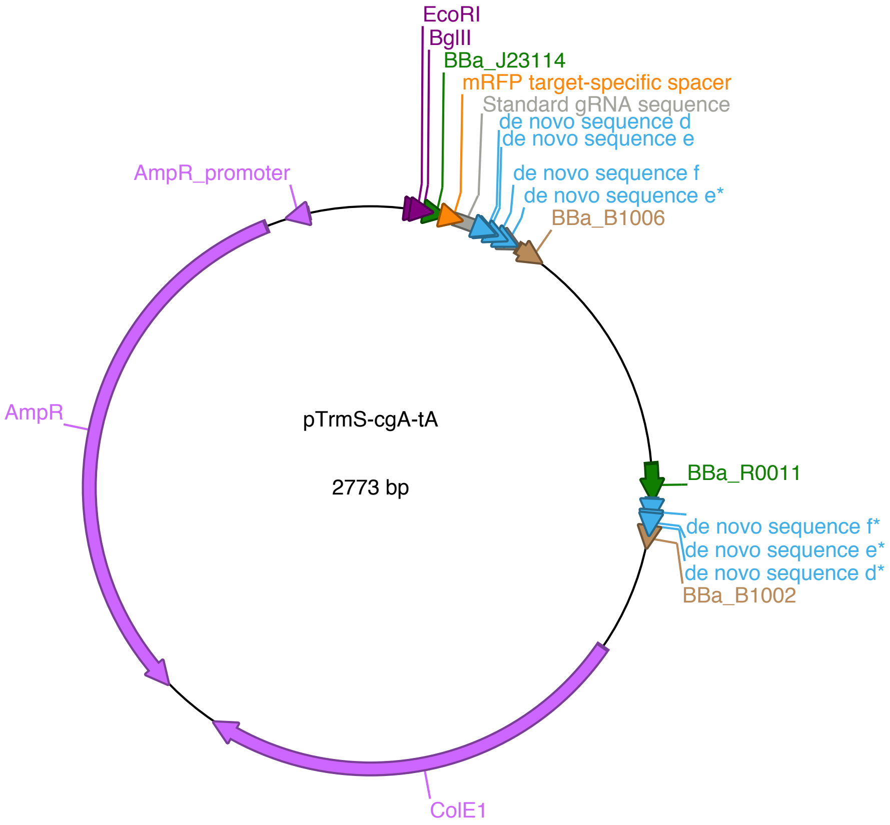


**Figure S8: Example plasmid map for terminator switch.** Plasmid: pTrmS-cgA-tA for cgRNA A + trigger A.

```
1 AATAGGCGTATCACGAGGCAGAATTTCAGATAAAAAAAATCCTTAGCTTTCGCTAAGGATGATTTCTGGAATTCTAAAGATCTTTTATGGCTAGCTCAGTCCTAGGTACAATGCTAGCAACTTTCAGTTTAGCGGTCTGTTTTAGAGCTAGAAATAGCAAGTTAAAATAAGGCTAGTCCGATCAAACGGGTAAACAAACA 200
  EcoRI  BglII  BBa_J23114  mRFP target-specific spacer  de novo sequence d
  Standard gRNA sequence  de novo sequence e
  de novo sequence

201 GGATAATTAAGGAGGCAGTACCCGGGCACCGAGTCGGTGCTTTTTTTAAAAAAAAACCCCGCCCCTGACAGGGCGGGGTTTTTTTGAAGCTTGGGCCCGAACAAAAACTCATCTCAGAAGAGGATCTGAATAGCGCCGTCGACCATCATCATCATCATCATTGAGTTTAAACGGTCTCCAGCTTGGCTGTTTTGGCGGA 400
  de novo sequence f  de novo sequence e*  BBa_B1006
  Standard gRNA sequence

401 TGAGAGAAGATTTTCAGCCTGATACAGATTAAATCAGAACGCAGAAGCGGTCTGATAAAACAGAATTTGCCTGGCGGCAGTAGCGCGGTGGTCCCACCTGACCCCATGCCGAACTCAGAAGTGAAACGCCGTAGCGCCGATGGTAGTGTGGGGTCTCCCCATGCGAGAGTAGGGAACTGCCAGGCATCAAATAAAACGAA 600

601 AGGCTCAGTCGAAAGACTGGGCCTTTCGTTTTATCTGTTGTTTGTCGGTGAACTGGATCAATTGTGAGCGGATAACAATTGACATTGTGAGCGGATAACAAGATACTGAGCACATACTGCCTCCTTAATTATCCTGTTTGTTTACCCGTTTGATCGCAAAAAACCCCGCTTCGGCGGGGTTTTTTCGCTCGAGTCTAGAC 800
  BBa_R0011  de novo sequence f*  de novo sequence e*
  de novo sequence d*
  BBa_B1002

801 TGCAGGCTTCCTCGCTCACTGACTCGCTGCGCTCGGTCGTTCGGCTGCGGCGAGCGGTATCAGCTCACTCAAAGGCGGTAATACGGTTATCCACAGAATCAGGGGATAACGCAGGAAAGAACATGTGAGCAAAAGGCCAGCAAAAGGCCAGGAACCGTAAAAAGGCCGCGTTGCTGGCGTTTTTCCATAGGCTCCGCCCC 1000
  ColE1

1001 CCTGACGAGCATCACAAAAATCGACGCTCAAGTCAGAGGTGGCGAAACCCGACAGGACTATAAAGATACCAGGCGTTTCCCCCTGGAAGCTCCCTCGTGCGCTCTCCTGTTCCGACCCTGCCGCTTACCGGATACCTGTCCGCCTTTCTCCCTTCGGGAAGCGTGGCGCTTTCTCATAGCTCACGCTGTAGGTATCTCAG 1200
  ColE1

1201 TTCGGTGTAGGTCGTTCGCTCCAAGCTGGGCTGTGTGCACGAACCCCCCGTTCAGCCCGACCGCTGCGCCTTATCCGGTAACTATCGTCTTGAGTCCAACCCGGTAAGACACGACTTATCGCCACTGGCAGCAGCCACTGGTAACAGGATTAGCAGAGCGAGGTATGTAGGCGGTGCTACAGAGTTCTTGAAGTGGTGGC 1400
  ColE1

1401 CTAACTACGGCTACACTAGAAGGACAGTATTTGGTATCTGCGCTCTGCTGAAGCCAGTTACCTTCGGAAAAAGAGTTGGTAGCTCTTGATCCGGCAAACAAACCACCGCTGGTAGCGGTGGTTTTTTTGTTTGCAAGCAGCAGATTACGCGCAGAAAAAAAGGATCTCAAGAAGATCCTTTGATCTTTTCTACGGGGTCT 1600
  ColE1

1601 GACGCTCAGTGGAACGAAAACTCACGTTAAGGGATTTTGGTCATGAGATTATCAAAAAGGATCTTCACCTAGATCCTTTTAAATTAAAAATGAAGTTTTAAATCAATCTAAAGTATATATGAGTAAACTTGGTCTGACAGTTACCAATGCTTAATCAGTGAGGCACCTATCTCAGCGATCTGTCTATTTCGTTCATCCAT 1800
  287
  ColE1  AmpR

1801 AGTTGCCTGACTCCCCGTCGTGTAGATAACTACGATACGGGAGGGCTTACCATCTGGCCCCAGTGCTGCAATGATACCGCGAGACCCACGCTCACCGGCTCCAGATTTATCAGCAATAAACCAGCCAGCCGGAAGGGCCGAGCGCAGAAGTGGTCCTGCAACTTTATCCGCCTCCATCCAGTCTATTAATTGTTGCCGGG 2000
  267
  AmpR

2001 AAGCTAGAGTAAGTAGTTCGCCAGTTAATAGTTTGCGCAACGTTGTTGCCATTGCTACAGGCATCGTGGTGTCACGCTCGTCGTTTGGTATGGCTTCATTCAGCTCCGGTTCCCAACGATCAAGGCGAGTTACATGATCCCCCATGTTGTGCAAAAAAGCGGTTAGCTCCTTCGGTCCTCCGATCGTTGTCAGAAGTAAG 2200
  200
  AmpR

2201 TTGGCCGCAGTGTTATCACTCATGGTTATGGCAGCACTGCATAATTCTCTTACTGTCATGCCATCCGTAAGATGCTTTTCTGTGACTGGTGAGTACTCAACCAAGTCATTCTGAGAATAGTGTATGCGGCGACCGAGTTGCTCTTGCCCGGCGTCAATACGGGATAATACCGCGCCACATAGCAGAACTTTAAAAGTGCT 2400
  133
  AmpR

2401 CATCATTGGAAAACGTTCTTCGGGGCGAAAACTCTCAAGGATCTTACCGCTGTTGAGATCCAGTTCGATGTAACCCACTCGTGCACCCAACTGATCTTCAGCATCTTTTACTTTCACCAGCGTTTCTGGGTGAGCAAAAACAGGAAGGCAAAATGCCGCAAAAAAGGGAATAAGGGCGACACGGAAATGTTGAATACTCA 2600
  67
  AmpR

2601 TACTCTTCCTTTTTCAATATTATTGAAGCATTTATCAGGGTTATTGTCTCATGAGCGGATACATATTTGAATGTATTTAGAAAAATAAACAAATAGGGGTTCCGCGCACATTTCCCCGAAAAGTGCCACCTGACGTCTAAGAAACCATTATTATCATGACATTAACCTATAAA 2773
  287
  AmpR  AmpR_promoter
```

**Figure S9: Example annotated plasmid sequence for terminator switch.** Plasmid: pTrmS-cgA-tA for cgRNA A + trigger A.

#### S3.2 Constitutively active splinted switch in *E. coli*

| Name | Parts | Figure | Legend |
|---|---|---|---|
| pdCas9-bacteria | Addgene plasmid #44249 | 3bc | |
| pg-NT | ColE1 ori; amp-R; BBa_J23114-NT_g-BBa_B1006 | 3b | No-target gRNA; Autofluorescence |
| | | 3c | Autofluorescence |
| pg-sfGFP | ColE1 ori; amp-R; BBa_J23114-sfGFP_g-BBa_B1006 | 3b | Standard gRNA |
| pSplS-cgA-nT | ColE1 ori; amp-R; BBa_J23108-SplS_cgA-BBa_B1006 | 3b | cgRNA |
| | | 3c | cgRNA A |
| pSplS-cgA-tA | ColE1 ori; amp-R; BBa_J23100-SplS_tA-BBa_B1006-BBa_B0015; BBa_J23108-SplS_cgA-BBa_B1006 | 3b | cgRNA + trigger |
| | See Figure S10, Figure S11 | 3c | cgRNA A, trigger $X_A$ |
| pSplS-cgA-tB | ColE1 ori; amp-R; BBa_J23100-SplS_tB-BBa_B1006-BBa_B0015; BBa_J23108-SplS_cgA-BBa_B1006 | 3c | cgRNA A, trigger $X_B$ |
| pSplS-cgA-tC | ColE1 ori; amp-R; BBa_J23100-SplS_tC-BBa_B1006-BBa_B0015; BBa_J23108-SplS_cgA-BBa_B1006 | 3c | cgRNA A, trigger $X_C$ |
| pSplS-cgB-nT | ColE1 ori; amp-R; BBa_J23108-SplS_cgB-BBa_B1006 | 3c | cgRNA B |
| pSplS-cgB-tA | ColE1 ori; amp-R; BBa_J23100-SplS_tA-BBa_B1006-BBa_B0015; BBa_J23108-SplS_cgB-BBa_B1006 | 3c | cgRNA B, trigger $X_A$ |
| pSplS-cgB-tB | ColE1 ori; amp-R; BBa_J23100-SplS_tB-BBa_B1006-BBa_B0015; BBa_J23108-SplS_cgB-BBa_B1006 | 3c | cgRNA B, trigger $X_B$ |
| pSplS-cgB-tC | ColE1 ori; amp-R; BBa_J23100-SplS_tC-BBa_B1006-BBa_B0015; BBa_J23108-SplS_cgB-BBa_B1006 | 3c | cgRNA B, trigger $X_C$ |
| pSplS-cgC-nT | ColE1 ori; amp-R; BBa_J23108-SplS_cgC-BBa_B1006 | 3c | cgRNA C |
| pSplS-cgC-tA | ColE1 ori; amp-R; BBa_J23100-SplS_tA-BBa_B1006-BBa_B0015; BBa_J23108-SplS_cgC-BBa_B1006 | 3c | cgRNA C, trigger $X_A$ |
| pSplS-cgC-tB | ColE1 ori; amp-R; BBa_J23100-SplS_tB-BBa_B1006-BBa_B0015; BBa_J23108-SplS_cgC-BBa_B1006 | 3c | cgRNA C, trigger $X_B$ |
| pSplS-cgC-tC | ColE1 ori; amp-R; BBa_J23100-SplS_tC-BBa_B1006-BBa_B0015; BBa_J23108-SplS_cgC-BBa_B1006 | 3c | cgRNA C, trigger $X_C$ |

**Table S7: Plasmids used with splinted switch cgRNAs in *E. coli*.**

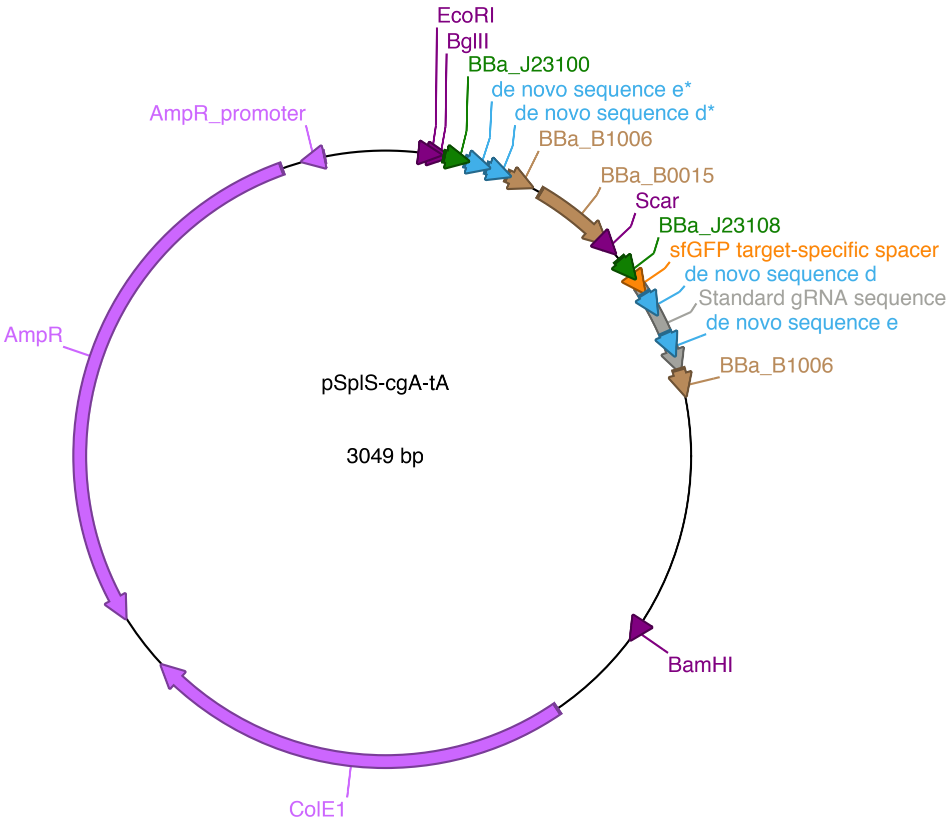


**Figure S10: Example plasmid map for splinted switch.** Plasmid: pSplS-cgA-tA for cgRNA A + trigger A.

```
1 AATAGGCGTATCACGAGGCAGAATTTCAGATAAAAAAAATCCTTAGCTTTCGCTAAGGATGATTTCTGGAATTCGTGGTAGGAGATCTGCATAAGGAGTTGACGGCTAGCTCAGTCCTAGGTACAGTGCTAGCAGGGTAAAGGAAGAGGAAGGTTTGTAAAGGGTGTTCTGGAAGGGACTTGGAATTGAACCGGCGTAAG 200
  EcoRI   BglII   BBa_J23100   de novo sequence e*   de novo sequence d*

201 GTGAAAAAAAAACCCCGCCCCTGACAGGGCGGGGTTTTTTTTCTACACCCTTCTAACAACTCCCAGGCATCAAATAAAACGAAAGGCTCAGTCGAAAGACTGGGCCTTTCGTTTTATCTGTTGTTTGTCGGTGAACGCTCTCTACTAGAGTCACACTGGCTCACCTTCGGGTGGGCCTTTCTGCGTTTATAGGATCTGAA 400
  de novo sequence d*   BBa_B0015   Scar
  BBa_B1006

401 ACATGACGAGCTGGTGAGCATAAGGAGCTGACAGCTAGCTCAGTCCTAGGTATAATGCTAGCCATCTAATTCAACAAGAATTGTTTTAGAGCTACACCTTACGCCGGTTCAATTCCAAGTCCCTTCCAGTAGCAAGTTAAAATAAGGCTAGTCCGTTATCAACTTAACACCCTTTACAAACCTTCCTCTTCCTTTACCCT 600
  BBa_J23108   sfGFP target-specific spacer   de novo sequence d   Standard gRNA sequence   de novo sequence e
  Standard gRNA sequence

601 AAGTGGCACCGAGTCGGTGCTTTTTTTAAAAAAAAACCCCGCCCCTGACAGGGCGGGGTTTTTTTTGCGCTATCAGTTGACACAGTGAAGCTTGGGCCCGAACAAAAACTCATCTCAGAAGAGGATCTGAATAGCGCCGTCGACCATCATCATCATCATCATTGAGTTTAAACGGTCTCCAGCTTGGCTGTTTTGGCGGA 800
  Standard gRNA sequence   BBa_B1006

801 TGAGAGAAGATTTTCAGCCTGATACAGATTAAATCAGAACGCAGAAGCGGTCTGATAAAACAGAATTTGCCTGGCGGCAGTAGCGCGGTGGTCCCACCTGACCCCATGCCGAACTCAGAAGTGAAACGCCGTAGCGCCGATGGTAGTGTGGGGTCTCCCCATGCGAGAGTAGGGAACTGCCAGGCATCAAATAAAACGAA 1000

1001 AGGCTCAGTCGAAAGACTGGGCCTTTCGTTTTATCTGTTGTTTGTCGGTGAACTGGATCCTTACTCGAGTCTAGACTGCAGGCTTCCTCGCTCACTGACTCGCTGCGCTCGGTCGTTCGGCTGCGGCGAGCGGTATCAGCTCACTCAAAGGCGGTAATACGGTTATCCACAGAATCAGGGGATAACGCAGGAAAGAACAT 1200
  BamHI

1201 GTGAGCAAAAGGCCAGCAAAAGGCCAGGAACCGTAAAAAGGCCGCGTTGCTGGCGTTTTTCCATAGGCTCCGCCCCCCTGACGAGCATCACAAAAATCGACGCTCAAGTCAGAGGTGGCGAAACCCGACAGGACTATAAAGATACCAGGCGTTTCCCCCTGGAAGCTCCCTCGTGCGCTCTCCTGTTCCGACCCTGCCGC 1400
  ColE1

1401 TTACCGGATACCTGTCCGCCTTTCTCCCTTCGGGAAGCGTGGCGCTTTCTCATAGCTCACGCTGTAGGTATCTCAGTTCGGTGTAGGTCGTTCGCTCCAAGCTGGGCTGTGTGCACGAACCCCCCGTTCAGCCCGACCGCTGCGCCTTATCCGGTAACTATCGTCTTGAGTCCAACCCGGTAAGACACGACTTATCGCCA 1600
  ColE1

1601 CTGGCAGCAGCCACTGGTAACAGGATTAGCAGAGCGAGGTATGTAGGCGGTGCTACAGAGTTCTTGAAGTGGTGGCCTAACTACGGCTACACTAGAAGGACAGTATTTGGTATCTGCGCTCTGCTGAAGCCAGTTACCTTCGGAAAAAGAGTTGGTAGCTCTTGATCCGGCAAACAAACCACCGCTGGTAGCGGTGGTTT 1800
  ColE1

1801 TTTTGTTTGCAAGCAGCAGATTACGCGCAGAAAAAAAGGATCTCAAGAAGATCCTTTGATCTTTTCTACGGGGTCTGACGCTCAGTGGAACGAAAACTCACGTTAAGGGATTTTGGTCATGAGATTATCAAAAAGGATCTTCACCTAGATCCTTTTAAATTAAAAATGAAGTTTTAAATCAATCTAAAGTATATATGAGT 2000
  ColE1

2001 AAACTTGGTCTGACAGTTACCAATGCTTAATCAGTGAGGCACCTATCTCAGCGATCTGTCTATTTCGTTCATCCATAGTTGCCTGACTCCCCGTCGTGTAGATAACTACGATACGGGAGGGCTTACCATCTGGCCCCAGTGCTGCAATGATACCGCGAGACCCACGCTCACCGGCTCCAGATTTATCAGCAATAAACCAG 2200
  287
  AmpR

2201 CCAGCCGGAAGGGCCGAGCGCAGAAGTGGTCCTGCAACTTTATCCGCCTCCATCCAGTCTATTAATTGTTGCCGGGAAGCTAGAGTAAGTAGTTCGCCAGTTAATAGTTTGCGCAACGTTGTTGCCATTGCTACAGGCATCGTGGTGTCACGCTCGTCGTTTGGTATGGCTTCATTCAGCTCCGGTTCCCAACGATCAAG 2400
  225
  AmpR

2401 GCGAGTTACATGATCCCCCATGTTGTGCAAAAAAGCGGTTAGCTCCTTCGGTCCTCCGATCGTTGTCAGAAGTAAGTTGGCCGCAGTGTTATCACTCATGGTTATGGCAGCACTGCATAATTCTCTTACTGTCATGCCATCCGTAAGATGCTTTTCTGTGACTGGTGAGTACTCAACCAAGTCATTCTGAGAATAGTGTA 2600
  159
  AmpR

2601 TGCGGCGACCGAGTTGCTCTTGCCCGGCGTCAATACGGGATAATACCGCGCCACATAGCAGAACTTTAAAAGTGCTCATCATTGGAAAACGTTCTTCGGGGCGAAAACTCTCAAGGATCTTACCGCTGTTGAGATCCAGTTCGATGTAACCCACTCGTGCACCCAACTGATCTTCAGCATCTTTTACTTTCACCAGCGTT 2800
  92
  AmpR

2801 TCTGGGTGAGCAAAAACAGGAAGGCAAAATGCCGCAAAAAAGGGAATAAGGGCGACACGGAAATGTTGAATACTCATACTCTTCCTTTTTCAATATTATTGAAGCATTTATCAGGGTTATTGTCTCATGAGCGGATACATATTTGAATGTATTTAGAAAAATAAACAAATAGGGGTTCCGCGCACATTTCCCCGAAAAGT 3000
  25
  AmpR   AmpR_promoter

3001 GCCACCTGACGTCTAAGAAACCATTATTATCATGACATTAACCTATAAA 3049
```

**Figure S11: Example annotated plasmid sequence for splinted switch.** Plasmid: pSplS-cgA-tA for cgRNA A + trigger A.

#### S3.3 Constitutively inactive toehold switch in *E. coli*

| Name | Parts | Figure | Legend |
|---|---|---|---|
| pdCas9-bacteria | Addgene plasmid #44249 | 4bc | |
| pg-NT | ColE1 ori; amp-R; BBa_J23114-NT_g-BBa_B1006 | 4b | No-target gRNA; Autofluorescence |
| | | 4c | Autofluorescence |
| pg-mRFP | ColE1 ori; amp-R; BBa_J23114-mRFP_g-BBa_B1006 | 4b | Standard gRNA |
| pToeS-cgA-nT | ColE1 ori; amp-R; BBa_J23100-BBa_B0050; BBa_J23108-ToeS_cgA | 4b | cgRNA |
| | | 4c | cgRNA A |
| pToeS-cgA-tA | ColE1 ori; amp-R; BBa_J23100-ToeS_tA-BBa_B0050; BBa_J23108-ToeS_cgA | 4b | cgRNA + trigger |
| | See Figure S12, Figure S13 | 4c | cgRNA A, trigger $X_A$ |
| pToeS-cgA-tB | ColE1 ori; amp-R; BBa_J23100-ToeS_tB-BBa_B0050; BBa_J23108-ToeS_cgA | 4c | cgRNA A, trigger $X_B$ |
| pToeS-cgA-tC | ColE1 ori; amp-R; BBa_J23100-ToeS_tC-BBa_B0050; BBa_J23108-ToeS_cgA | 4c | cgRNA A, trigger $X_C$ |
| pToeS-cgB-nT | ColE1 ori; amp-R; BBa_J23100-BBa_B0050; BBa_J23108-ToeS_cgB | 4c | cgRNA B |
| pToeS-cgB-tA | ColE1 ori; amp-R; BBa_J23100-ToeS_tA-BBa_B0050; BBa_J23108-ToeS_cgB | 4c | cgRNA B, trigger $X_A$ |
| pToeS-cgB-tB | ColE1 ori; amp-R; BBa_J23100-ToeS_tB-BBa_B0050; BBa_J23108-ToeS_cgB | 4c | cgRNA B, trigger $X_B$ |
| pToeS-cgB-tC | ColE1 ori; amp-R; BBa_J23100-ToeS_tC-BBa_B0050; BBa_J23108-ToeS_cgB | 4c | cgRNA B, trigger $X_C$ |
| pToeS-cgC-nT | ColE1 ori; amp-R; BBa_J23100-BBa_B0050; BBa_J23108-ToeS_cgC | 4c | cgRNA C |
| pToeS-cgC-tA | ColE1 ori; amp-R; BBa_J23100-ToeS_tA-BBa_B0050; BBa_J23108-ToeS_cgC | 4c | cgRNA C, trigger $X_A$ |
| pToeS-cgC-tB | ColE1 ori; amp-R; BBa_J23100-ToeS_tB-BBa_B0050; BBa_J23108-ToeS_cgC | 4c | cgRNA C, trigger $X_B$ |
| pToeS-cgC-tC | ColE1 ori; amp-R; BBa_J23100-ToeS_tC-BBa_B0050; BBa_J23108-ToeS_cgC | 4c | cgRNA C, trigger $X_C$ |

**Table S8:** Plasmids used with toehold switch cgRNAs in *E. coli*

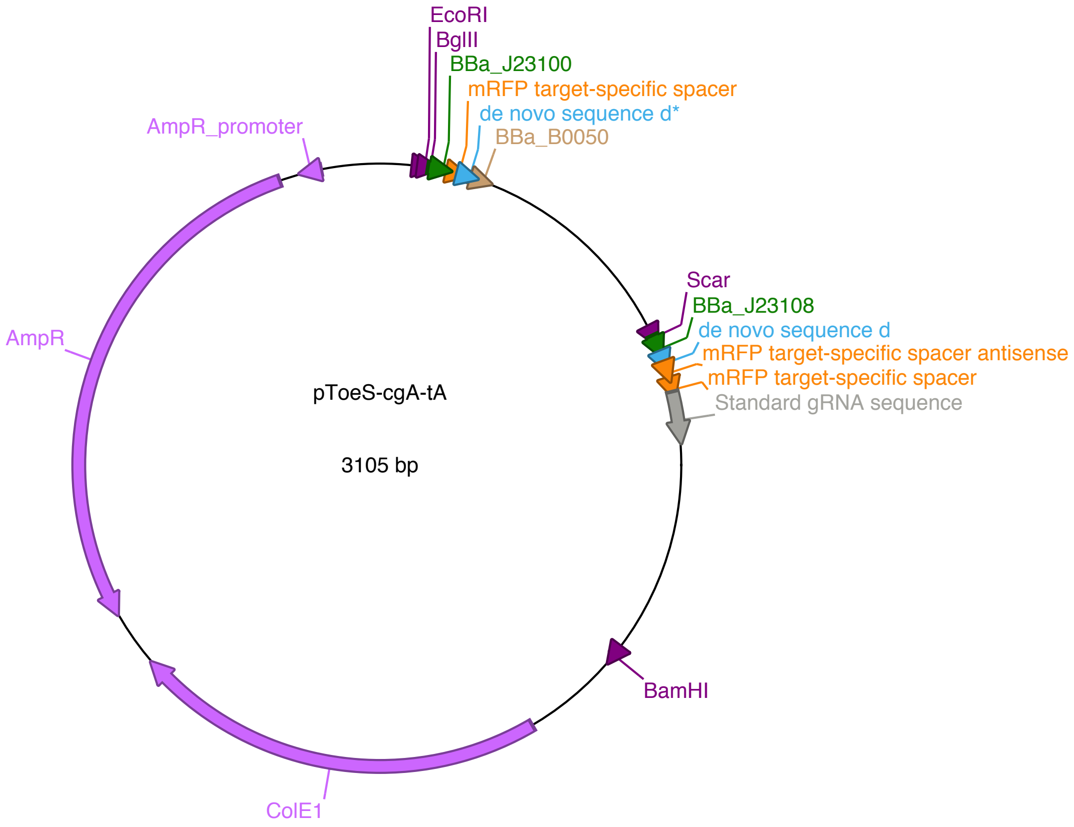


**Figure S12: Example plasmid map for toehold switch.** Plasmid: pToeS-cgA-tA for cgRNA A + trigger A.

1 AATAGGCGTATCACGAGGCAGAATTTCAGATAAAAAAAATCCTTAGCTTTCGCTAAGGATGATTTCTGGAATTCTAAAGATCTTTGACGGCTAGCTCAGTCCTAGGTACAGTGCTAGCAACTTTCAGTTTAGCGGTCTTAATACAACGAACATAAAAAAAGGATCTCAAGAAGATCCTTTGATTTTTGAAGCTTGGGCCC 200

EcoRI BglII BBa_J23100 mRFP target-specific spacer BBa_B0050

de novo sequence d*

201 GAACAAAAACTCATCTCAGAAGAGGATCTGAATAGCGCCGTCGACCATCATCACCATCATCATTGAGTTTAAACGGTCTCCAGCTTGGCTGTTTTGGCGGATGAGAGAAGATTTTCAGCCTGATACAGATTAAATCAGAACGCAGAAGCGGTCTGATAAAACAGAATTTGCCTGGCGGCAGTAGCGCGGTGGTCCCACCT 400

401 GACCCCATGCCGAACTCAGAAGTGAAACGCCGTAGCGCCGATGGTAGTGTGGGGTCTCCCCATGCGAGAGTAGGGAACTGCCAGGCATCAAATAAAACGAAAGGCTCAGTCGAAAGACTGGGCCTTTCGTTTTATCTGTTGTTTGTCGGTGAACTGGATCTCTGACAGCTAGCTCAGTCCTAGGTATAATGCTAGCATGT 600

Scar BBa_J23108 de novo sequence d

601 TCGTTGTATTAAGACCGCTAAACTGAAAGTTACACGCCCAACTTTCAGTTTAGCGGTCTGTTTTAGAGCTAGAAATAGCAAGTTAAAATAAGGCTAGTCCGTTATCAACTTGAAAAAGTGGCACCGAGTCGGTGCTTTTTTTGAAGCTTGGGCCCGAACAAAAACTCATCTCAGAAGAGGATCTGAATAGCGCCGTCGAC 800

de novo sequence d mRFP target-specific spacer

mRFP target-specific spacer antisense Standard gRNA sequence

801 CATCATCATCATCATCATTGAGTTTAAACGGTCTCCAGCTTGGCTGTTTTGGCGGATGAGAGAAGATTTTCAGCCTGATACAGATTAAATCAGAACGCAGAAGCGGTCTGATAAAACAGAATTTGCCTGGCGGCAGTAGCGCGGTGGTCCCACCTGACCCCATGCCGAACTCAGAAGTGAAACGCCGTAGCGCCGATGGT 1000

1001 AGTGTGGGGTCTCCCCATGCGAGAGTAGGGAACTGCCAGGCATCAAATAAAACGAAAGGCTCAGTCGAAAGACTGGGCCTTTCGTTTTATCTGTTGTTTGTCGGTGAACTGGATCCTTACTCGAGTCTAGACTGCAGGCTTCCTCGCTCACTGACTCGCTGCGCTCGGTCGTTCGGCTGCGGCGAGCGGTATCAGCTCAC 1200

BamHI

1201 TCAAAGGCGGTAATACGGTTATCCACAGAATCAGGGGATAACGCAGGAAAGAACATGTGAGCAAAAGGCCAGCAAAAGGCCAGGAACCGTAAAAAGGCCGCGTTGCTGGCGTTTTTCCATAGGCTCCGCCCCCCTGACGAGCATCACAAAAATCGACGCTCAAGTCAGAGGTGGCGAAACCCGACAGGACTATAAAGATA 1400

ColE1

1401 CCAGGCGTTTCCCCCTGGAAGCTCCCTCGTGCGCTCTCCTGTTCCGACCCTGCCGCTTACCGGATACCTGTCCGCCTTTCTCCCTTCGGGAAGCGTGGCGCTTTCTCATAGCTCACGCTGTAGGTATCTCAGTTCGGTGTAGGTCGTTCGCTCCAAGCTGGGCTGTGTGCACGAACCCCCCGTTCAGCCCGACCGCTGCG 1600

ColE1

1601 CCTTATCCGGTAACTATCGTCTTGAGTCCAACCCGGTAAGACACGACTTATCGCCACTGGCAGCAGCCACTGGTAACAGGATTAGCAGAGCGAGGTATGTAGGCGGTGCTACAGAGTTCTTGAAGTGGTGGCCTAACTACGGCTACACTAGAAGGACAGTATTTGGTATCTGCGCTCTGCTGAAGCCAGTTACCTTCGGA 1800

ColE1

1801 AAAAGAGTTGGTAGCTCTTGATCCGGCAAACAAACCACCGCTGGTAGCGGTGGTTTTTTTGTTTGCAAGCAGCAGATTACGCGCAGAAAAAAAGGATCTCAAGAAGATCCTTTGATCTTTTCTACGGGGTCTGACGCTCAGTGGAACGAAAACTCACGTTAAGGGATTTTGGTCATGAGATTATCAAAAAGGATCTTCAC 2000

ColE1

2001 CTAGATCCTTTTAAATTAAAAATGAAGTTTTAAATCAATCTAAAGTATATATGAGTAAACTTGGTCTGACAGTTACCAATGCTTAATCAGTGAGGCACCTATCTCAGCGATCTGTCTATTTCGTTCATCCATAGTTGCCTGACTCCCCGTCGTGTAGATAACTACGATACGGGAGGGCTTACCATCTGGCCCCAGTGCTG 2200

287

AmpR

2201 CAATGATACCGCGAGACCCACGCTCACCGGCTCCAGATTTATCAGCAATAAACCAGCCAGCCGGAAGGGCCGAGCGCAGAAGTGGTCCTGCAACTTTATCCGCCTCCATCCAGTCTATTAATTGTTGCCGGGAAGCTAGAGTAAGTAGTTCGCCAGTTAATAGTTTGCGCAACGTTGTTGCCATTGCTACAGGCATCGTG 2400

244

AmpR

2401 GTGTCACGCTCGTCGTTTGGTATGGCTTCATTCAGCTCCGGTTCCCAACGATCAAGGCGAGTTACATGATCCCCCATGTTGTGCAAAAAAGCGGTTAGCTCCTTCGGTCCTCCGATCGTTGTCAGAAGTAAGTTGGCCGCAGTGTTATCACTCATGGTTATGGCAGCACTGCATAATTCTCTTACTGTCATGCCATCCGT 2600

177

AmpR

2601 AAGATGCTTTTCTGTGACTGGTGAGTACTCAACCAAGTCATTCTGAGAATAGTGTATGCGGCGACCGAGTTGCTCTTGCCCGGCGTCAATACGGGATAATACCGCGCCACATAGCAGAACTTTAAAAGTGCTCATCATTGGAAAACGTTCTTCGGGGCGAAAACTCTCAAGGATCTTACCGCTGTTGAGATCCAGTTCGA 2800

111

AmpR

2801 TGTAACCCACTCGTGCACCCAACTGATCTTCAGCATCTTTTACTTTCACCAGCGTTTCTGGGTGAGCAAAAACAGGAAGGCAAAATGCCGCAAAAAAGGGAATAAGGGCGACACGGAAATGTTGAATACTCATACTCTTCCTTTTTCAATATTATTGAAGCATTTATCAGGGTTATTGTCTCATGAGCGGATACATATTT 3000

44

AmpR AmpR_promoter

3001 GAATGTATTTAGAAAAATAAACAAATAGGGGTTCCGCGCACATTTCCCCGAAAAGTGCCACCTGACGTCTAAGAAACCATTATTATCATGACATTAACCTATAAA 3105

AmpR_promoter

**Figure S13: Example annotated plasmid sequence for toehold switch.** Plasmid: pToeS-cgA-tA for cgRNA A + trigger A.

#### S3.4 Plasmid for expression of lacI + dCas9

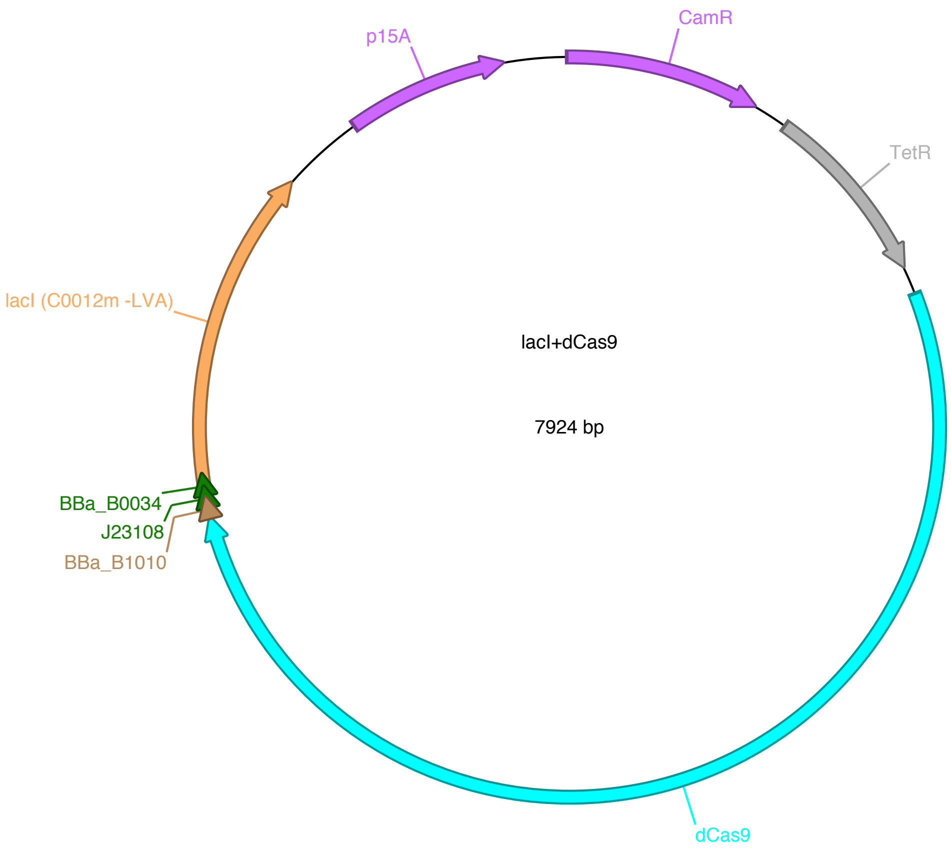


**Figure S14: Plasmid map for pdCas9+lacI**

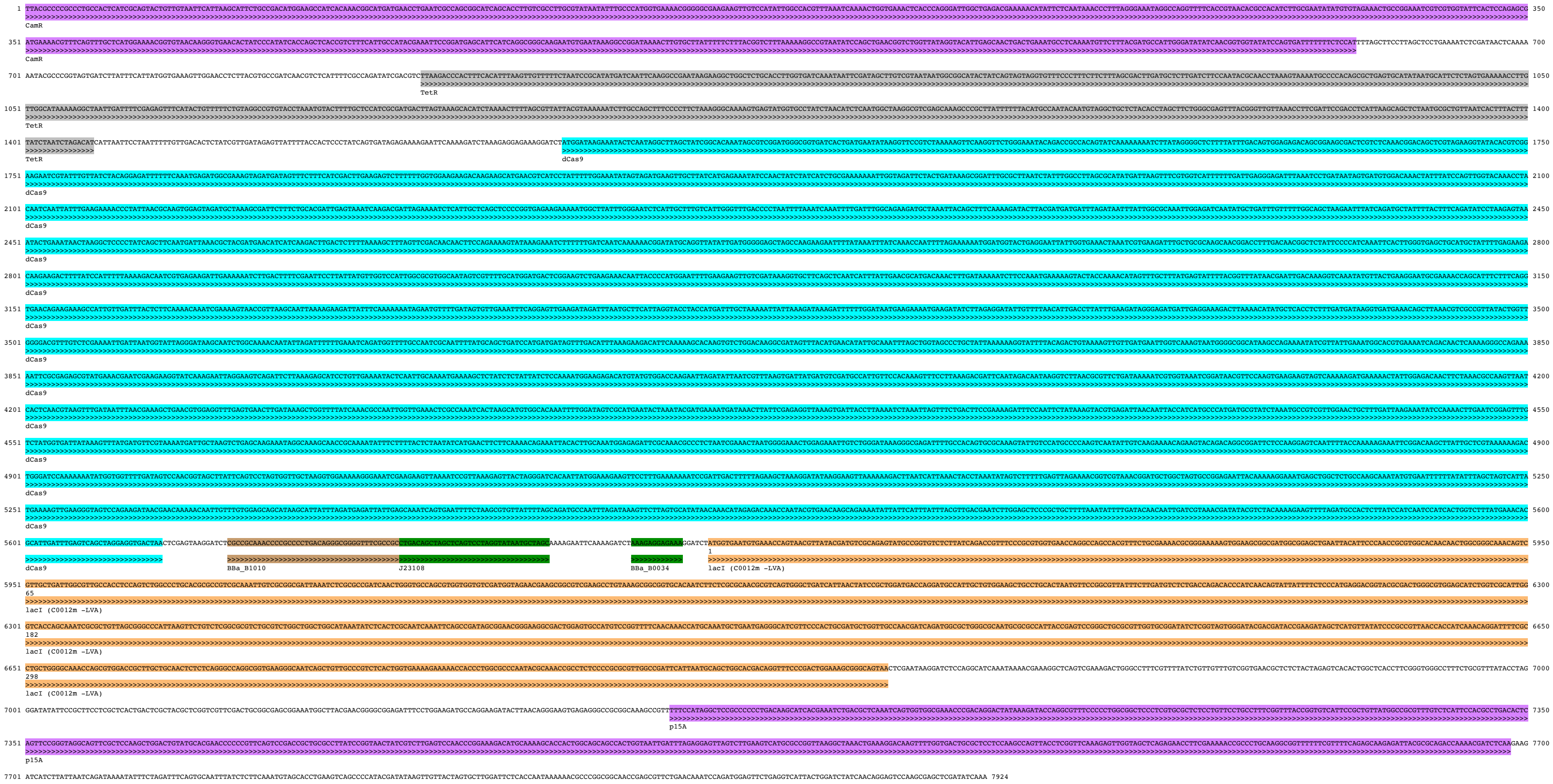


**Figure S15: Annotated plasmid sequence for expression of lacI + dCas9.** Plasmid: pdCas9+lacI.

### S4 Schematics of putative ON and OFF states

#### S4.1 Constitutively active terminator switch cgRNA

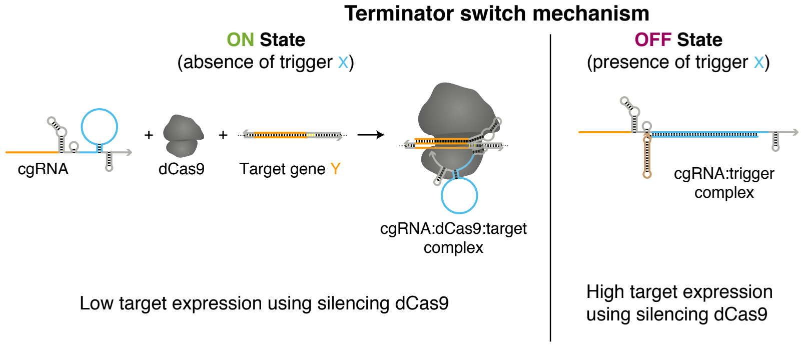


**Figure S16:** Schematics of putative ON and OFF states for the terminator switch mechanism. ON state: the terminator switch cgRNA is constitutively active, directing the function of protein effector dCas9 to a target gene Y in the absence of trigger; the extended loop and modified sequence domains in the terminator region (blue) are intended not to interfere with the activity of the cgRNA:dCas9 complex. OFF state: in the presence of RNA trigger X, hybridization of the trigger is intended to form a structure incompatible with cgRNA mediation of dCas9 function.

#### S4.2 Constitutively active splinted switch cgRNA

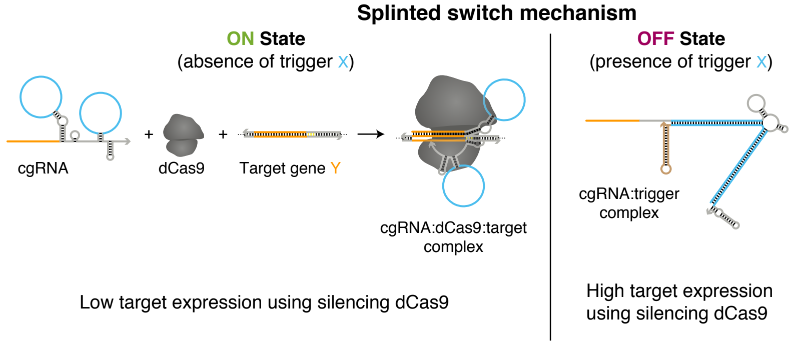


**Figure S17:** Schematics of putative ON and OFF states for the splinted switch mechanism. ON state: the splinted switch cgRNA is constitutively active, directing the function of protein effector dCas9 to a target gene Y in the absence of trigger; the extended loops in the Cas9 handle and terminator region (blue) are intended not to interfere with the activity of the cgRNA:dCas9 complex. OFF state: in the presence of RNA trigger X, hybridization of the trigger is intended to form a splint that is structurally incompatible with cgRNA mediation of dCas9 function.

#### S4.3 Constitutively inactive toehold switch cgRNA

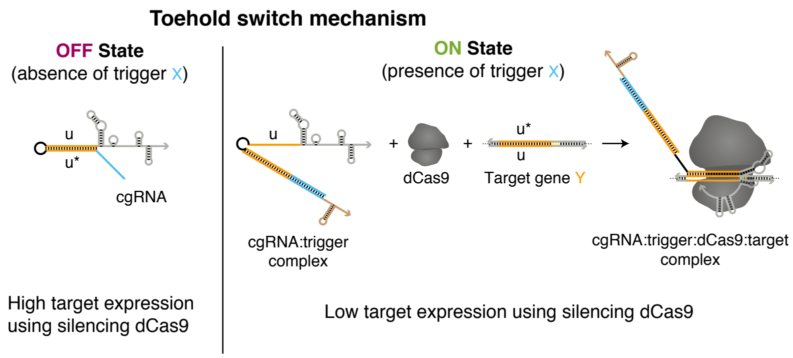


**Figure S18:** Schematics of putative OFF and ON states for the toehold switch mechanism. OFF state: The toehold switch cgRNA is constitutively inactive; the target-binding region (domain “u”; orange) is initially sequestered by a 5’ extension to inhibit recognition of target gene Y. ON state: in the presence of RNA trigger X, hybridization of the trigger to this extension via the toehold region (blue) is intended to de-sequester the target-binding region and enable cgRNA direction of dCas9 function to target gene Y.

### S5 Flow cytometry replicates

#### S5.1 Constitutively active terminator switch in *E. coli*

##### S5.1.1 ON state, OFF state, and conditional response (cf. Figure 2b)

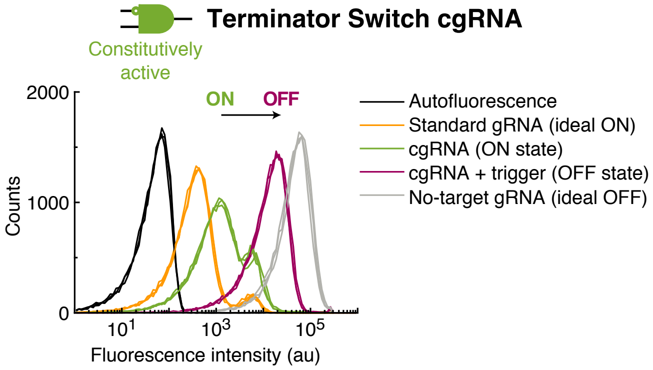


**Figure S19: Flow cytometry replicates for terminator switch ON state, OFF state, and conditional response in *E. coli* (cf. Figure 2b).** Single-cell fluorescence intensities. Expression of RNA trigger X toggles the cgRNA from ON→OFF, leading to an increase in fluorescence. Induced expression (aTc) of silencing dCas9 and constitutive expression of mRFP target gene Y and either: standard gRNA (ideal ON state), cgRNA(ON state), cgRNA + RNA trigger X (OFF state; trigger expression is IPTG-induced), no-target gRNA that lacks target-binding region (ideal OFF state). Autofluorescence: cells with no mRFP. Traces of the same color correspond to $N = 3$ replicate wells assayed on the same day (20,000 cells per well).

##### S5.1.2 Orthogonal library studies (cf. Figure 2c)

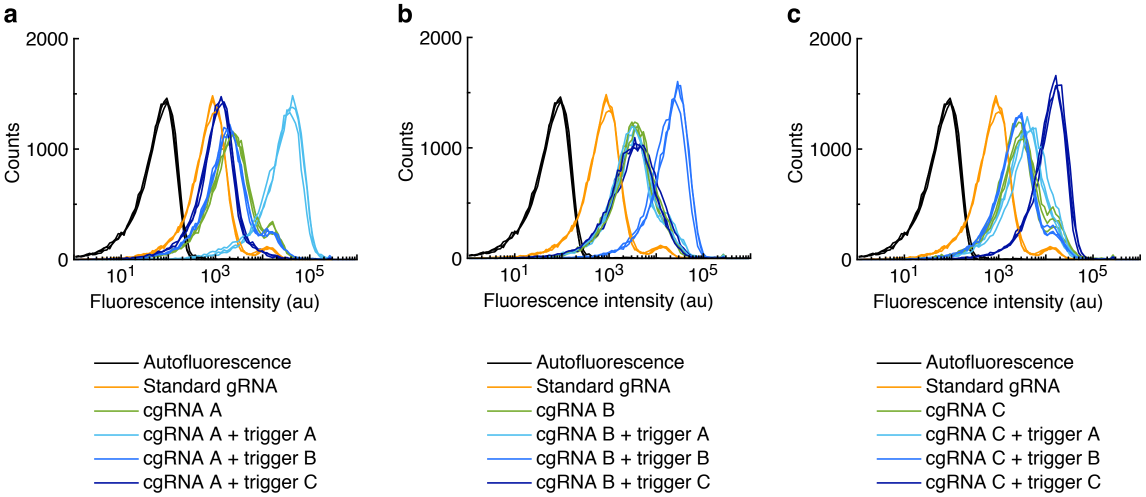


**Figure S20: Flow cytometry replicates for terminator switch orthogonal response in *E. coli* (cf. Figure 2c).** (a) cgRNA A. (b) cgRNA B. (c) cgRNA C. Single-cell fluorescence intensities. Induced expression (aTc) of silencing dCas9 and constitutive expression of mRFP target gene Y and either: standard gRNA, cgRNA without trigger, cgRNA + cognate trigger, or cgRNA + a non-cognate trigger (trigger expression is IPTG-induced). Autofluorescence: cells with no mRFP. Expression of the cognate RNA trigger ($X_A$ for cgRNA A, $X_B$ for cgRNA B, $X_C$ for cgRNA C) toggles the cgRNA from ON→OFF, leading to an increase in fluorescence. Traces of the same color correspond to $N = 3$ replicate wells assayed on the same day (20,000 cells per well).

#### S5.2 Constitutively active splinted switch in *E. coli*

##### S5.2.1 ON state, OFF state, and conditional response (cf. Figure 3b)

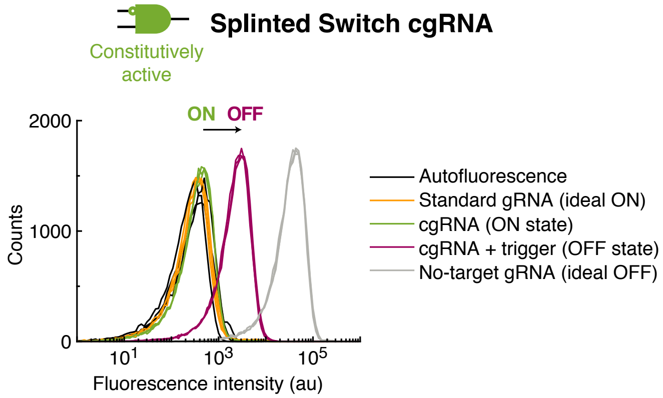


**Figure S21: Flow cytometry replicates for splinted switch ON state, OFF state, and conditional response in *E. coli* (cf. Figure 3b).** Single-cell fluorescence intensities. Expression of RNA trigger X toggles the cgRNA from ON→OFF, leading to an increase in fluorescence. Induced expression (aTc) of silencing dCas9 and constitutive expression of sfGFP target gene Y and either: standard gRNA (ideal ON state), cgRNA(ON state), cgRNA + RNA trigger X (OFF state), no-target gRNA that lacks target-binding region (ideal OFF state). Autofluorescence: cells with no sfGFP. Traces of the same color correspond to $N = 3$ replicate wells assayed on the same day (20,000 cells per well).

##### S5.2.2 Orthogonal library studies (cf. Figure 3c)

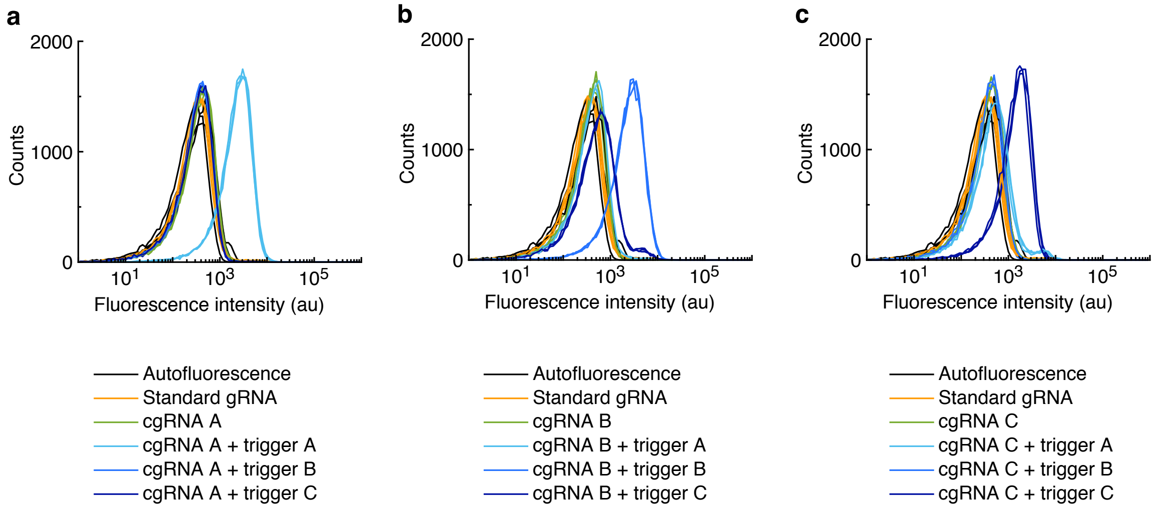


**Figure S22: Flow cytometry replicates for splinted switch orthogonal response in *E. coli* (cf. Figure 3c).** (a) cgRNA A. (b) cgRNA B. (c) cgRNA C. Single-cell fluorescence intensities. Induced expression (aTc) of silencing dCas9 and constitutive expression of sfGFP target gene Y and either: standard gRNA, cgRNA without trigger, cgRNA + cognate trigger, or cgRNA + a non-cognate trigger. Autofluorescence: cells with no sfGFP. Expression of the cognate RNA trigger ($X_A$ for cgRNA A, $X_B$ for cgRNA B, $X_C$ for cgRNA C) toggles the cgRNA from ON→OFF, leading to an increase in fluorescence. Traces of the same color correspond to $N = 3$ replicate wells assayed on the same day (20,000 cells per well).

#### S5.3 Constitutively inactive toehold switch in *E. coli*

##### S5.3.1 ON state, OFF state, and conditional response (cf. Figure 4b)

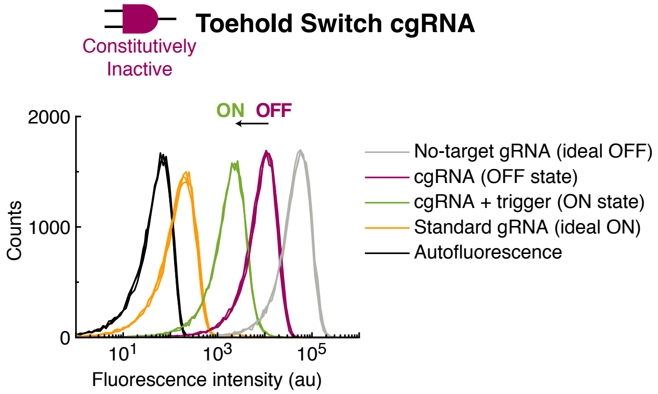


**Figure S23: Flow cytometry replicates for toehold switch ON state, OFF state, and conditional response in *E. coli* (cf. Figure 4b).** Single-cell fluorescence intensities. Expression of RNA trigger X toggles the cgRNA from OFF→ON, leading to a decrease in fluorescence. Induced expression (aTc) of silencing dCas9 and constitutive expression of mRFP target gene Y and either: no-target gRNA that lacks target-binding region (ideal OFF state), cgRNA (OFF state), cgRNA + RNA trigger X (ON state), standard gRNA (ideal ON state). Autofluorescence: cells with no mRFP. Traces of the same color correspond to $N = 3$ replicate wells assayed on the same day (20,000 cells per well).

##### S5.3.2 Orthogonal library studies (cf. Figure 4c)

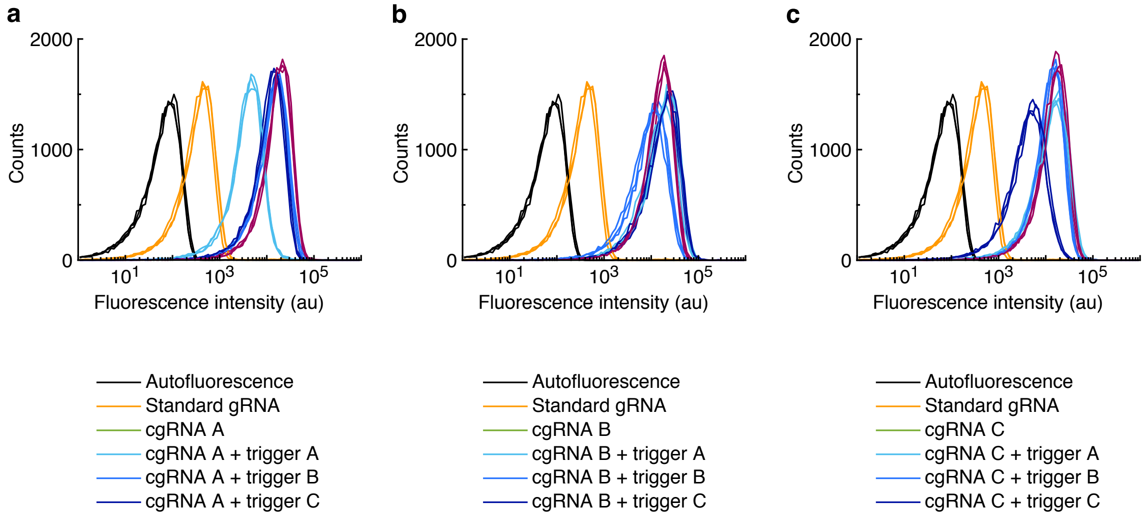


**Figure S24: Flow cytometry replicates for toehold switch orthogonal response in *E. coli* (cf. Figure 4c).** Flow cytometry fluorescence assay shows selective activation of Toehold Switch cgRNA with expression of cognate RNA trigger. (a) cgRNA A. (b) cgRNA B. (c) cgRNA C. Single-cell fluorescence intensities. Induced expression (aTc) of silencing dCas9 and constitutive expression of mRFP target gene Y and either: standard gRNA, cgRNA without trigger, cgRNA + cognate trigger, or cgRNA + a non-cognate trigger. Autofluorescence: cells with no mRFP. Expression of the cognate RNA trigger ($X_A$ for cgRNA A, $X_B$ for cgRNA B, $X_C$ for cgRNA C) toggles the cgRNA from OFF→ON, leading to decrease in fluorescence. Traces of the same color correspond to $N = 3$ replicate wells assayed on the same day (20,000 cells per well).
